# CELLoGeNe - an Energy Landscape Framework for Logical Networks Controlling Cell Decisions

**DOI:** 10.1101/2022.02.09.479734

**Authors:** Emil Andersson, Mattias Sjö, Keisuke Kaji, Victor Olariu

**Affiliations:** Computational Biology and Biological Physics, Department of Astronomy and Theoretical Physics, Lund University, Lund, Sweden; Centre for Regenerative Medicine, University of Edinburgh, Edinburgh, UK

## Abstract

Experimental and computational efforts are constantly made to elucidate mechanisms controlling cell fate decisions during development and cell reprogramming. One powerful method is to consider cell commitment and reprogramming as movements in an energy landscape. Here, we develop CELLoGeNe (Computation of Energy Landscapes of Logical Gene Networks), which maps Boolean implementation of gene regulatory networks (GRNs) into energy landscapes. CELLoGeNe removes inadvertent symmetries in the energy landscapes normally arising from standard Boolean operators. Furthermore, CELLoGeNe provides a tool for visualising multi-dimensional energy landscapes and a platform to stochastically probe and analyse the shapes of the computed landscapes corresponding to the epigenetic landscapes for development and reprogramming. We demonstrate CELLoGeNe on a GRN governing maintenance and self-renewal of pluripotency, identifying attractors experimentally validated. We also apply CELLoGeNe on a network controlling reprogramming from mouse embryonic fibroblast (MEF) to induced pluripotent stem cells (iPSCs) where we identify potential roadblocks as attractors. CELLoGeNe is a general framework that can be applied to various biological systems offering a broad picture of intracellular dynamics otherwise inaccessible with existing methods.

## Introduction

The human body contains more than 200 different types of cells, which is a small number compared to the approximately 10^13^ cells in total (***Alberts et al., 2008***). The distinguishing feature between different cell types is which genes are activated, or *expressed*. In natural development, cells can only transition from pluripotent cells into more specialised cells, a process governed by changing gene expressions. However, experimental efforts have been made to reprogram cells, from specialised cells into pluripotent cells or other specialised cell types directly, for instance (***Davis et al., 1987; Xie et al., 2004; Takahashi and Yamanaka, 2006; Takahashi et al., 2007; Zhou et al., 2008; Aasen et al., 2008; Kim et al., 2009; Loh et al., 2010; Szabo et al., 2010; Staerk et al., 2010; Vierbuchen et al., 2010; Ieda et al., 2010; Huang et al., 2011***). Cell reprogramming is conducted by forcing expression of transcription factors (TFs), for recent overviews see (***Aydin and Mazzoni, 2019; Wang et al., 2021***). In particular, Takahashi and Yamanaka showed that it is possible to reprogram adult fibroblasts to induced pluripotent cells (iPSCs) by forcing the expression of just four TFs (Oct4, Klf4, Sox2, c-Myc) in both mouse and human cells (***Takahashi and Yamanaka, 2006; Takahashi et al., 2007***). However, conversion rates are poor with many roadblocks in the reprogramming paths (***O’Malley et al., 2013; Chantzoura et al., 2015***). Many computational efforts have been made to elucidate the intricate gene regulatory networks (GRNs) governing cell decisions and reprogramming, for example (***Reinitz et al., 1995; Gardner et al., 2000; Chen et al., 2000; Chickarmane et al., 2012; Dunn et al., 2014; Xu et al., 2014; Olariu et al., 2016, 2017a***,b; ***Dunn et al., 2019***).

The human genome is estimated to contain at least 30 000 genes (***Roest Crollius et al., 2000***), which either can be expressed at various levels or not expressed at all. Taking the whole genome with a continuous expression level into account when constructing a model for reprogramming is not feasible due to the vast space of possible gene expression patterns a cell can exhibit. Fortunately, there are quite a few simplifications that can be done, which still yield good results. In a model, it is often possible to only consider a handful up to to a few dozens of genes shown to play an important role in the reprogramming process, depending on the complexity of the proposed model. However, the gene expression space still has an infinite amount of points, leading to another common simplification, namely, binarising the gene expression into either being OFF (0) or ON (1). This was first suggested by Kauffman already in the ‘60s (***Kauffman, 1969***). With this simplification, the gene expression space is reduced to a finite space with 2^*N*^ states, where *N* is the number of genes used in the model. Such Boolean models have been previously explored, see (***Mendoza et al., 1999; Fauré et al., 2006; Davidson, 2010; Peter et al., 2012***).

Considering cell commitment and reprogramming as movements in an energy landscape is a powerful tool for analysing decisions, which has been used in several previous studies (***Bhattacharya et al., 2011; Wang et al., 2011; Zhou et al., 2012; Mojtahedi et al., 2016; Olariu et al., 2017a; Corson and Siggia, 2017; Sáez et al., 2021***). Energy landscapes have their roots in a qualitative metaphor for cell fate decisions where Waddington envisaged pluripotent cells as marbles at the top of a hill, while cell differentiation was represented by them rolling down different available paths on the valleys and eventually stopping at a specialised state (***Waddington, 1957***). Energy landscapes are based on the same conceptual idea, but with a quantitative mapping from GRNs. Each theoretically possible cell state is assigned a specific energy value depending on the genes’ activation statuses and how they are regulated according to the GRN. In physics, energy is a quantity that is as small as possible when a system is completely relaxed. Hence, low energies are awarded to cell states where the network is “comfortable”, i.e. where the gene expressions and regulatory forces agree, and conversely for high energies. In this paradigm, cell states correspond to the minima in the landscape, i.e. attractors in the landscape where the metaphoric marbles come to a stop. Cell fate decisions are, hence, represented by the attractors’ *basins of attraction*, which include all those states from which the marbles inevitably will roll down into the attractors; when a marble has entered a basin it cannot escape.

Here, we develop CELLoGeNe (<u>C</u>omputation of <u>E</u>nergy <u>L</u>andscapes of <u>Lo</u>gical <u>Ge</u>ne <u>Ne</u>tworks), which maps Boolean implementation of gene regulatory networks into energy landscapes. Applying standard Boolean rules is accompanied by symmetries that may not be desired. CELLoGeNe solves this issue by implementing a three-state extension to Boolean logic where a gene’s expression (ON or OFF) is decoupled from its effect on a target gene (positive (+1), negative (−1), or neutral (0), see Methods-Logical representation and Methods-Three-state logic.

An energy landscape is a function with as many dimensions as the number of genes (*N*) considered in the GRN. This leads to visualisation challenges. CELLoGeNe provides a tool to plot multidimensional discrete energy landscapes (up to seven dimensions). This is achieved by considering each binary cell state as a corner on an N-dimensional hypercube, which is flattened out to two dimensions.

In order to analyse the strengths of the basins of attractions, we implemented a stochastic method that probes the shape of the energy landscape through weighted random walk. In essence, we release a large number of the metaphorical marbles at each cell state of the energy landscape, add a noise level, and record at which state they stop. This yields the probability of reaching the different attractors from each state of the landscape. By specifically analysing how marbles may move from one attractor to another when noise is included, one effectively analyses cell development and reprogramming.

We applied CELLoGeNe to a GRN governing maintenance and self-renewal of pluripotency (***Dunn et al., 2014***). Here, we identify attractors that are experimentally validated (***Dunn et al., 2014***), establishing that energy landscapes computed by CELLoGeNe indeed can correspond to experimental findings. Thereafter, we applied CELLoGeNe to a GRN controlling reprogramming from mouse embryonic fibroblast (MEF) to iPSCs. The topology of the network was extracted from experimental and computational results published by our laboratories and others. We identified known cell states as attractors along with a few additional attractors that predict the existence of potential bottlenecks in cell reprogramming.

CELLoGeNe is a general framework that can be applied to various biological systems controlled by a regulatory network. An important strength of CELLoGeNe is that it provides a broad picture of intracellular dynamics, as opposed to existing methods like solving systems of rate equations where it is hard to find all potential stable states. In fact, CELLoGeNe is not confined only to biology applications as it can be applied to any system that is governed by a logical regulatory network.

## Results

The state of the art of computationally studying cell reprogramming consists of two modelling strategies: (i) dynamical systems approach - constructing rate equations for change in gene expression and numerically solve them finding stable states, (ii) Boolean approach - binarising the GRN and logically update the network states until a stable state is reached (***Olariu and Peterson, 2019***). These two approaches suffer from a drawback, namely, they do not provide an overview of all available stable states connected to a GRN. This drawback can be overcome by calculating free energy landscapes linked to the GRN. To this end, we developed CELLoGeNe, a method for translating Boolean models for GRNs into free energy landscapes. CELLoGeNe uses a three-valued logic which provides a better model of gene interactions when translated to the energy picture.

### Overview of CELLoGeNe

The structural workflow of CELLoGeNe is illustrated in ***Figure 1***. In order to apply CELLoGeNe to a problem experimental gene expression data and a GRN topology are needed. The data must be binarised into gene expression ON (1) or OFF (0). The GRN may either be previously known, or hypothetical topologies can be tested through CELLoGeNe and validated by experimental data. After the GRN topology is fixed the logical combinations of operators combining multiple input signals at the GRN nodes are identified (***Figure 1***, Methods-Combining input signals with operators). This can either be done manually (by expert curation), or by letting CELLoGeNe test possible configurations, either exhaustively or stochastically (Methods-Testing configurations of operators). After the operators have been assigned, the energy landscape is calculated (Methods-The discrete energy). The attractors in the energy landscape are compared to the ones from binarised experimental data. If the data and energy landscape does not agree, the search through possible combinations of logical operators is continued. However, if the search was exhaustive, then the GRN topology does not agree with the experimental data and should be further optimised. If the data and energy landscape agree, it means that the landscape could be biologically plausible and can be further analysed (***Figure 1***). The analysis includes finding all minima in the energy landscape which provides predictions for cell states which could play a role as barriers in cell transitions between experimentally identified attractors. CELLoGeNe also provides a stochastic method for analysing the basins of attraction for each landscape minimum (Methods-Marble simulations). Moreover, we developed a visualisation tool, enabling us to depict energy landscapes with more than three dimensions (Methods-Visualisation of high-dimensional energy landscapes).

**Figure 1.**
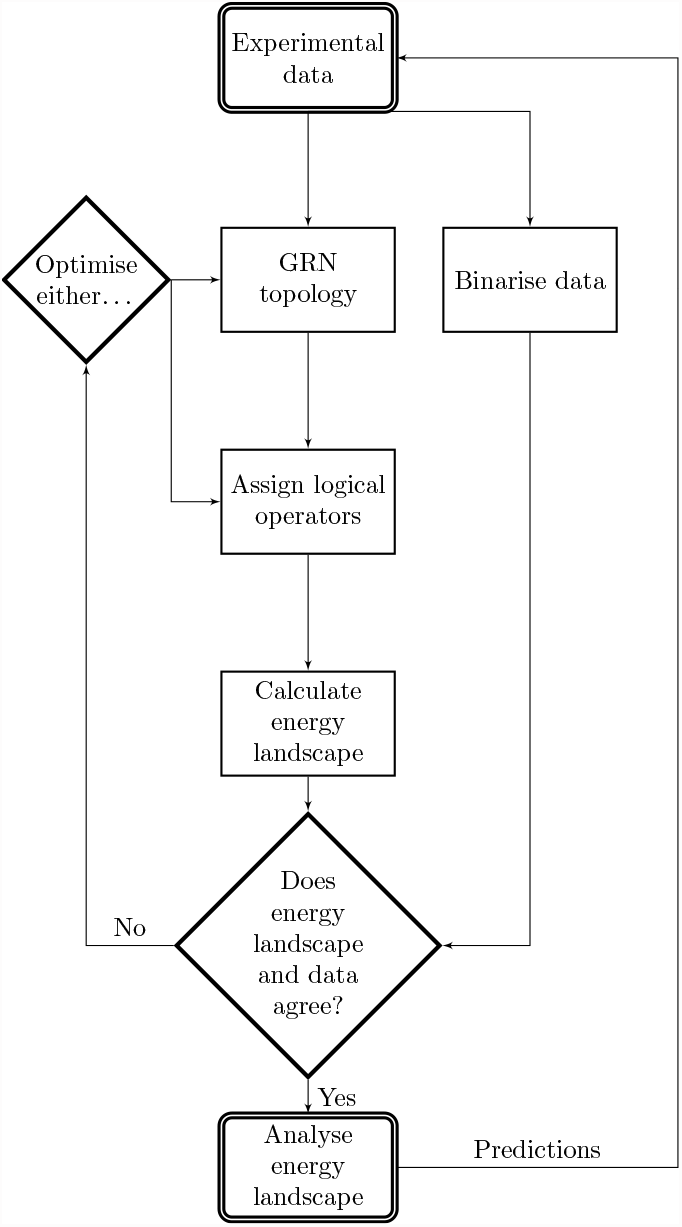
Flowchart illustrating the main workflow of CELLoGeNe.

### Demonstration on a toy model

In the following section, we demonstrate CELLoGeNe by applying it to a toy model. We consider the GRN in ***Figure 2A***, with five genes (nodes) A to E with (**→**) depicting activation and 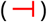 repression. A five-gene GRN results in 2^5^ = 32 possible cell states. We use CELLoGeNe to compute an energy landscape. Low energy (−1) is assigned to a cell state when the input signal at each node and their expression values agree while high energy (+1) is assigned in case of disagreement. In case of no input signal, we assign neutral energy (0).

**Figure 2.**
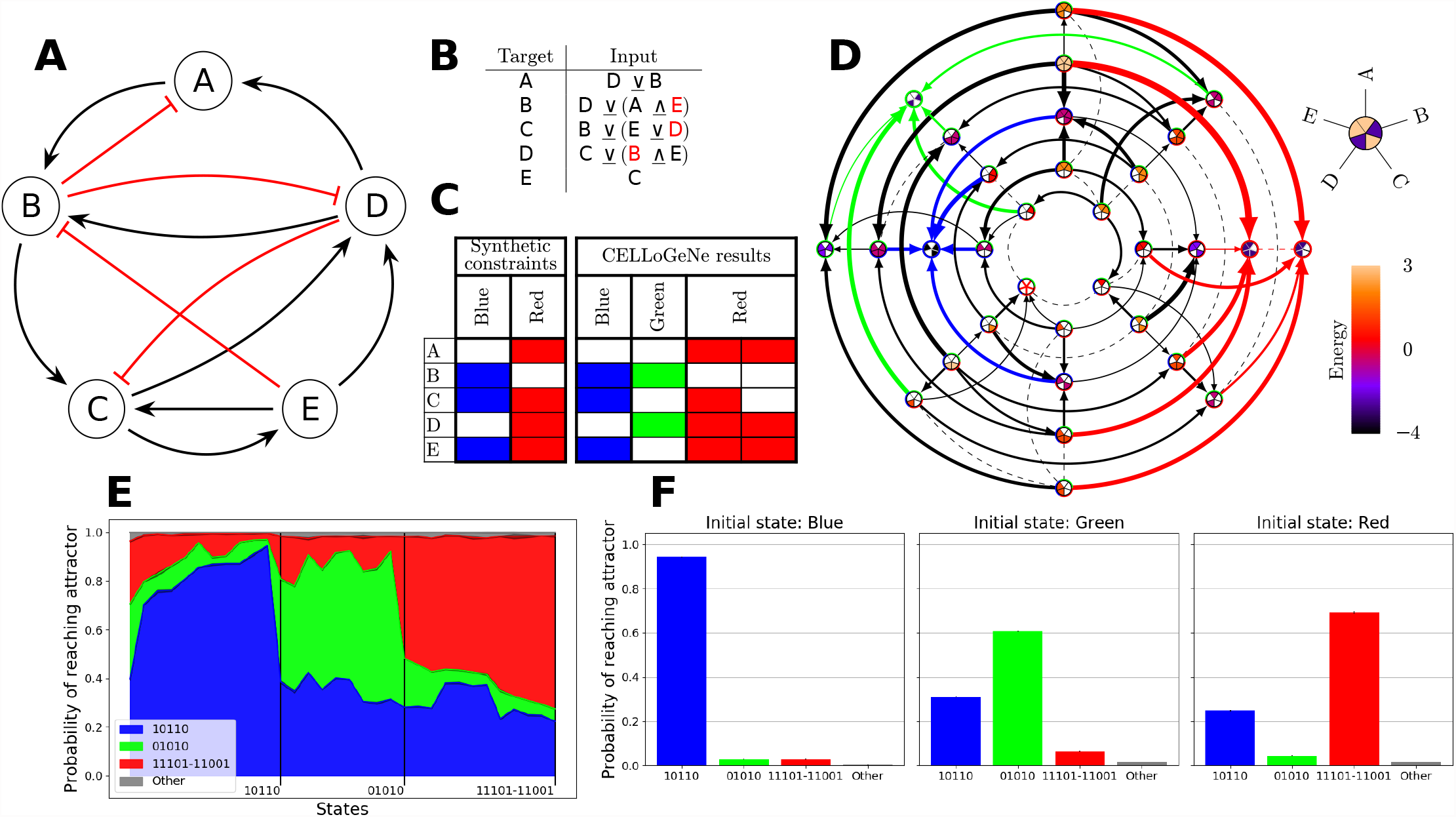
Application of CELLoGeNe on a toy model. (**A**) A GRN for a simple 5 gene toy model. (**B**) Example of a configuration of logical operators fulfilling the synthetic constraints. (**C**) The synthetic constraints and resulting attractors for the configuration in **B**. (**D**) The energy landscape corresponding to the configuration of logical operators displayed in **B**. Each landscape node represents a possible state of the logical cell where the colour of the node shows the energy value of that state, given by the colour bar to the right. Each node is divided into five sectors, where each sector corresponds to the five genes as indicated by the legend above the colour bar. Filled sectors correspond expressed genes while white sectors correspond to unexpressed genes. The arrows connect neighbouring states and point towards lower energies. The thickness of the arrows depends on the magnitude of the energy gradient. Dashed lines illustrate flat segments in the landscape, i.e. the two neighbouring states having the same energy value. The arrows to the three attractors in the landscape are colour coded. The coloured ring around each landscape node shows in which basins of attraction the node resides. (**E**) Overview of the basin sizes and relative strengths given from marble simulations where 10 000 marbles were initialised in each state with noise level *β* = 1.0. (**F**) Simulated reprogramming with marble simulations using the same settings as in **E**.

As outlined above, the two prerequisites for applying CELLoGeNe is to have a GRN and experimental data; however, since this is a toy model, we do not have any gene expression data. Hence, we introduced synthetic constraints by considering the states (A = OFF, B = ON, C = ON, D = OFF, E = ON) and (A = ON, B = OFF, C = ON, D = ON, E = ON) as attractors, depicted as blue and red in ***Figure 2C***. The next step is to assign logical operators to combine the input signals. For instance, gene C receives input both from gene B and E (***Figure 2A***), thus the dual signal must be combined with an operator into a single signal (Methods-Combining input signals with operators). For simplicity, we only considered the two operators <u>∨</u> and <u>∧</u> as defined in Methods-Combining input signals with operators. CELLoGeNe constructed the energy landscape for each possible configuration of these two operators (Methods-The discrete energy), and we picked one valid configuration which yielded an energy landscape fulfilling the synthetic constraints (***Figure 2B***). The full energy landscape is displayed in ***Figure 2D*** and the found attractors are summarised in ***Figure 2C***. Each landscape node represents a cell state, and each cell state is denoted by a pie chart. If the piece of pie corresponding to e.g. gene A (see the legend in ***Figure 2D***) is filled with a colour, it means that gene A is expressed. The colour of the node reflects the energy value of that cell state, with energy levels given by the colour bar to the right in ***Figure 2D***. Neighbouring states are connected with arrows, where their thickness is proportional to the magnitude of the energy difference between those states. Dashed lines illustrate neighbouring states with the same energy, i.e. flat areas of the energy landscape. For more details on the construction of landscape plots, see Methods-Visualisation of high-dimensional energy landscapes.

In addition to the attractors given by the synthetic constraints (blue and red), we identified another attractor in the landscape (green). Since the green state is not previously known, CELLoGeNe predicts that a stable state with only genes B and D ON should exist for this system. Additionally, the red state has a neighbouring state with the same energy value, i.e. the state is degenerate. A degenerate state suggests that the gene which can be either ON or OFF, in this case gene C, does not affect that specific cell type. Since the blue attractor has lower energy than both the red and green attractors, it presumably is a stronger attractor. Note that energy −5 (−*N* in general) means that all the genes’ expressions in the GRN agree with their input signal and +5 (+*N*) means that all disagree.

The coloured circle around each landscape node (***Figure 2D***) shows which attractors are possible to reach from the particular cell state by following the direction of the arrows or dashed lines, i.e. the basins of attraction. To investigate the basins’ relative strengths, we developed a stochastic method to analyse the probability of reaching each attractor from each cell state (Methods-Marble simulations). In principle, we let a metaphorical marble perform a weighted random walk in the landscape. At each step, the marble either rolls in one random direction, with a probability given by the difference in energy of the neighbouring states, or does not move. We introduced a noise level so that the marble, with a small probability, also can roll uphill. If the marble comes to a stop and stays at the same state for 3 updates, we consider the marble to have reached a final state. By repeating such simulations many times, the probability of reaching each attractor from a specific initial cell state is obtained. These stochastic simulations with noise allow all attractors to be reached from each cell state, although, with different probabilities. The result from these simulations is presented in ***Figure 2E***. On the horizontal axis, we consider initial states, while the vertical axis shows the probability of reaching the different attractors. The plot is divided into three sections corresponding to different dominating attractors, e.g. the blue attractor is dominating in the left section. This plot provides a qualitative illustration of the strength of each attractor. At the top of ***Figure 2E***, there is a small grey band, which means that for a small fraction of simulations, the marble stopped at other states than any of the three attractors. This is due to noise in the simulations.

In cell reprogramming, a cell transitions from one stable cell type to another. In our framework, this corresponds to the marble moving from one landscape attractor to another. The noise level perturbs the marble out from the attractor, which then either rolls back or away into another attractor (***Figure 2F***). In this landscape, the blue attractor corresponds to a cell state which is more difficult to reprogram from. The green and red attractors represent cell states amenable to be reprogrammed towards the blue cell type.

### Maintenance of naïve pluripotency

In this section, we apply CELLoGeNe to a GRN governing self-renewal and maintenance of naïve pluripotency in mouse embryonic stem (ES) cells. *In vitro*, mouse ES cells can self-renew indefinitely, without losing their differentiation capacity into all three germ layers, in certain culture conditions including medium containing leukaemia inhibitory factor (LIF), inhibitor of glycogen synthase kinase 3 (CHIR99021, CH), and inhibitor of mitogen-activated protein (PD0325901, PD), where the latter two are referred as two inhibitors (2i) (***Ying et al., 2008***). A minimal GRN controlling maintenance of mouse ES cells in four combinations of media LIF + 2i, 2i, LIF + CH, LIF + PD was put forward by (***Dunn et al., 2014***). Here, we apply CELLoGeNe to the published minimal GRN and use the available binarised expression data for each stable state for the four combinations of medium components (***Figure 3A*** and ***Figure 3C***).

**Figure 3.**
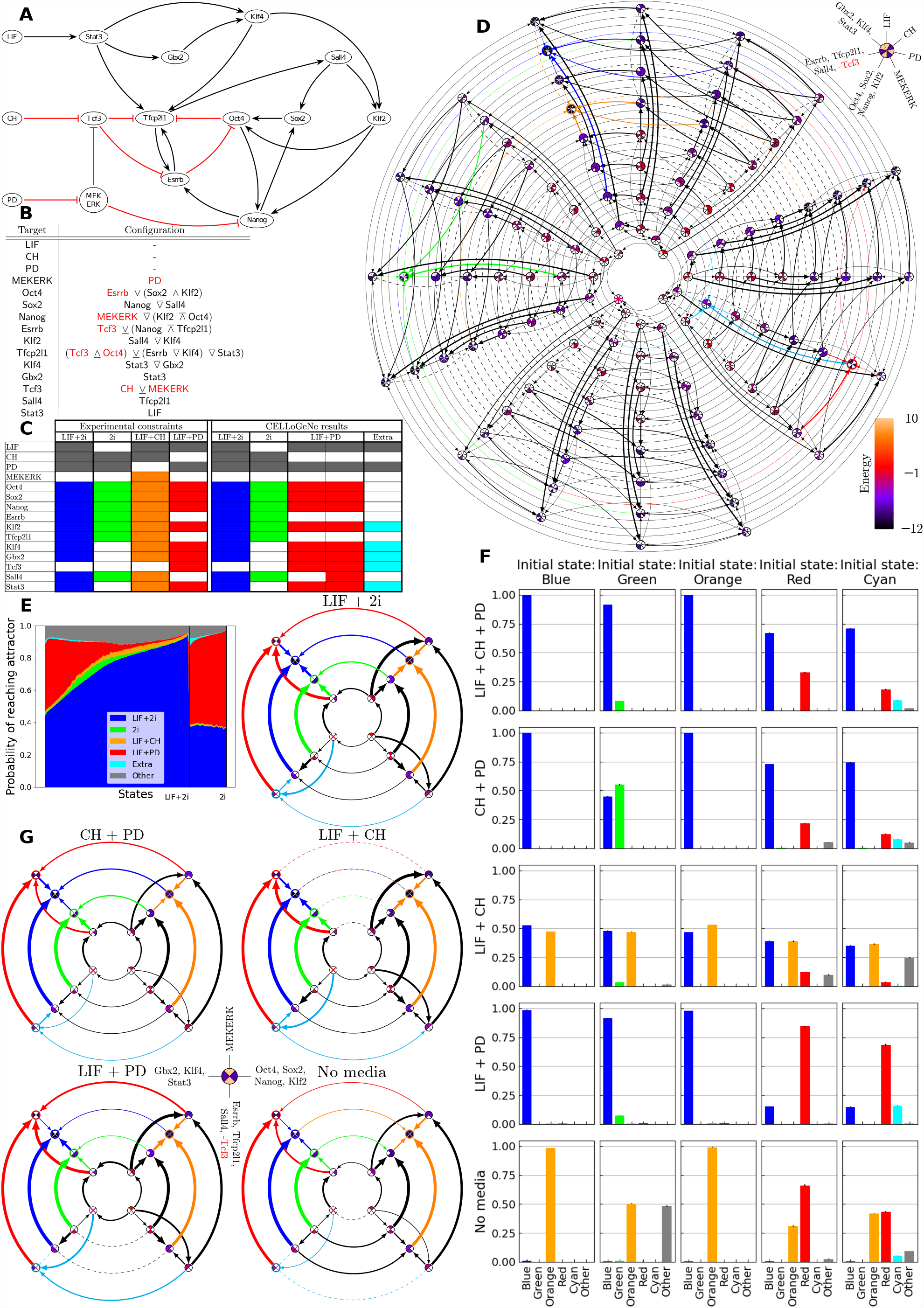
(**A**) GRN describing maintenance of pluripotency (***Dunn et al., 2014***). Black arrows represent activation, while blunted red arrows represent repression. (**B**) Chosen configuration of logical operators. (**C**) Experimental constraints (***Dunn et al., 2014***) and the resulting attractors from CELLoGeNe. (**D**) Energy landscape where groups of genes have been collected into the same nodes (see legend in the top-right). Each landscape node represents a possible state of the logical cell where the colour of the node shows the energy value of that state, where the values are given by the colour bar to the right. Each node is divided into seven sectors, where the sectors correspond to the genes as indicated in the legend. Filled sectors correspond to expressed genes while white sectors correspond to unexpressed genes. The arrows connect neighbouring states and points toward lower energies. The thickness of the arrows is a proportional to the magnitude of the energy gradient. Dashed lines illustrate flat segments in the landscape, i.e. two neighbouring states having the same energy value. The connections to the attractors of the landscapes are colour coded. (**E**) Overview of the relative basin sizes and strengths given from stochastic marble simulations. 100 marbles per initial state were simulated with noise-level *β* = 1.5. (**F**) Simulated differentiation. The distribution of final attractors of marbles initialised from the attractors with constant media using *β* = 1.5 and 10 000 marbles per initial state. (**G**) The five disconnected energy landscapes illustrating differentiation of cells transferred between media. **Figure 3–Figure supplement 1**. Number of times the 10 most common attractors and experimental constraints occur in the 10^6^ calculated energy landscapes. **Figure 3–Figure supplement 2**. The 3 configurations with the lowest degeneracy out of the 163 valid configurations found when trying 10^6^ configurations.

The GRN (***Figure 3A***) consists of 15 nodes: 12 TFs and 3 medium components acting as input nodes. In the CELLoGeNe framework, all of these are treated equally and are referred to as genes. In this case, there are 2^15^ = 32 768 possible cell states, which is considerably more than the 32 in the toy model. Furthermore, since most of the GRN nodes have multiple inputs, there are more than 28 trillions configurations of operators^1^ when using all 6 of the proposed operators (Methods-Configurations of operators).

With this amount of possible configurations, it is not computationally feasible to exhaustively test all configurations; hence, we performed exhaustive searches for cases when we considered only three sets of two operators 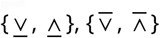 and {↑, ↓}. Moreover, we stochastically tried 10^6^ configurations with all operators. No configurations from the three two-operator sets yielded energy landscapes where all four experimental constraints (***Figure 3C***) were fulfilled. In the stochastic search, the state in LIF + 2i was the most prevalent attractor occurring in 46 % of the 10^6^ calculated energy landscapes (***Figure 3 – Figure Supplement 1***). Thus, the GRN seems extremely robust when it comes to maintaining pluripotency in LIF + 2i. Also the stable state in LIF + CH was one of the 10 most prevalent attractors, occurring in 23 % of the landscapes. In total, 163 valid configurations were found. Three of them (***Figure 3 – Figure Supplement 2***) were deemed more plausible than the others since they had lower degeneracy in the attractors. However, none of these configurations were biologically satisfactory when the operator combinations were scrutinised. Firstly, all three configurations required that both CH and MEKERK are ON in order to repress Tcf3. This is not reasonable since the presence of PD inhibits MEKERK, therefore leaving CH without any effect on Tcf3, which is against the known behaviour of CH. Secondly, in some instances, repressive input is effectively used as activation. An example of this can be seen in the input for Oct4 in Configuration 2: Oct4 ← (Sox2 ↓ Klf 2) 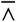 Esrrb. Here, Oct4 can only receive an activation signal if all three input genes are ON, even the repressive Esrrb. There is no way to receive repression of Oct4 from Esrrb with this configuration. Thirdly, in some cases, an inhibitor and an activator are combined with an <u>∧</u> -operator, i.e. both the activator and inhibitor must be ON in order for the target gene to receive a repressive signal. An example of this can be seen in Configuration 3 for Tfcp2𝗅1 ← ((Stat3 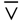 (Esrrb <u>∧</u> Oct4)) <u>∧</u> Klf 4) <u>∨</u> Tcf3, where both the activating Esrrb and repressive Oct4 must be ON for Tf cp2I1 to be repressed by Oct4, effectively turning Esrrb into a repressor.

To avoid unreasonable configurations, we manually curated a configuration (***Figure 3B***), where none of the repressors effectively worked as activators or needed an activator in order to repress its target, and where CH had the authority to inhibit Tcf3 on its own. The resulting attractors are presented in ***Figure 3C***. Note that the experimentally stable state in LIF + CH (orange) is not an attractor in the energy landscape. Nonetheless, we do not consider this to be a problem. The GRN (***Figure 3A***) cannot produce the orange state as a single attractor while still following the reasoning about repressors above since it is very close to the stable state in LIF + 2i (blue). The only non-media gene that differ between the experimentally stable states blue and orange is MEKERK (MEKERK = OFF in blue and MEKERK = ON in orange). Thus, when PD = OFF, then MEKERK is free and can be either ON or OFF with the same energy, which corresponds to the same state as blue except in another medium. This is clearly illustrated in the energy landscape plot (***Figure 3D***).^2^ Here, it is clear that the orange state in LIF + CH and the blue state in LIF + 2i only differ by two genes, and the landscape is flat for changing MEKERK (orange dashed line in the direction of 11 o’clock). Then changing medium from PD = OFF to PD = ON is favourable in energy but can not happen spontaneously in experiments.

The manually curated configuration predicts an additional stable state, labelled “Extra” with cyan colour in ***Figure 3***. This state is in LIF + PD medium but is not a pluripotent state. Only Klf 2 of the pluripotent genes^3^ is ON. However, this attractor is very weak as shown by the marble simulations (***Figure 3E***). The strongest attractor is the blue state in LIF + 2i media, which can be reached from all initial states and is the dominating attractor for approximately 80 % of the states. The red attractor in LIF + PD is the dominating attractor for around 20 % of the states. The orange LIF + CH state is fairly even, but with a low probability of being reached from all initial states. The green 2i attractor, on the other hand, is quite weak and can practically only be reached from about half of the states, suggesting that it is easier to maintain the pluripotency with LIF present. The fact that the blue attractor is found to be the strongest one is in good agreement with the results from (***Dunn et al., 2014***) where it was shown that ES cells cultured in LIF + 2i are remarkably robust. Over the whole domain, there is a 5 % to 15 % probability of reaching other states than the attractors. This probably is a sign that large regions of the energy landscapes are flat. This might correspond to spontaneous exit from pluripotency towards somatic or other unidentified cell fates.

Finally, we performed simulations representing transferring a cell colony cultured in one medium to another as done experimentally in (***Dunn et al., 2014***). This is done with the marble simulations, but with constraints prohibiting the marble to roll in a direction that changes the medium. The simulations take place on different, disconnected, parts of the energy landscape (***Figure 3G***), which can be thought of as an *in silicio* version of putting cell cultures in a Petri dish containing a specific medium *in vitro*. Simulations were performed for each of the 25 possible combinations of initial attractors and media (including absence of medium). The resulting attractor distributions are displayed in ***Figure 3F*** where the medium is the same in each row, and the initial state is the same in each column. This model simulation predicted that the blue state is the strongest attractor, even in other media than LIF + 2i. The reason for this can be seen in the energy landscapes with disconnected media (***Figure 3G***). Blue is the sole minimum in three of the media, and a degenerate minimum together with orange in LIF + CH. It is only in the landscape with no medium where blue is not a minimum, being replaced by orange.

Merging groups of genes for visualisation purpose has the disadvantage that the resulting landscape plot is not an exact representation of the energy landscape as the neighbouring nodes in the merged landscape are not neighbouring cell states^4^. For this reason, the green state, for instance, does not appear as a minimum in the CH + PD landscape. From the green state, there is lower energy in the neighbouring state where Gbx2, Klf 4 and Stat3 are all ON; however, in the full landscape, a neighbouring state only changes one gene’s expression. Even though these merged landscapes do not show connections between direct neighbours, they are still very useful for an illustrative purpose as they show main directions in the landscape, which is important in the stochastic marble simulations.

The disconnected landscapes (***Figure 3G***) explain the attractor distributions. For instance, the green attractor is stronger for the green initial state in CH + PD than in LIF + 2i. Comparing the corresponding landscapes, there is a larger energy difference between the green and blue state in LIF + 2i than in CH + PD, illustrated both by the colour scale and the thickness of the arrow, yielding higher probability for cells to move from the green state to the blue. The situation is similar for the red and blue states in LIF + PD compared to the other media. The orange and blue state are degenerate in LIF + CH (***Figure 3G*** - blue and orange dotted line), which corresponds to the third row of ***Figure 3F*** where reaching the blue and orange states has the same probability, with a slight bias towards the initial state. The orange state is the strongest attractor state when simulating the no-medium scenario. Experimentally, only MEKERK should be ON in the absence of media (***Dunn et al., 2014***). However, the presence of the orange state as an attractor in the absence of media can be explained. In CELLoGeNe, when no medium components are present, then both MEKERK and Stat3 receive a neutral input signal. Then, it is clearly beneficial for MEKERK to be ON and Tcf3 to be OFF since that agree with MEKERK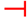Tcf3. If all the other genes are OFF (i.e. the cell state with only MEKERK = ON), then only Tcf3 and Nanog receive input signal and the cell state has energy −2. The local landscape area around this state is largely flat since many of the genes have no input signal. Turning on Stat3 would in itself not yield lower energy, however, the state with all downstream targets from Stat3 turned ON would be beneficial, which corresponds to the orange state. It should be noted that for the green initial state with in no medium, CELLoGeNe identified a large fraction of “Other” states. This is probably due to the green state already having many genes OFF, thus being close to the large flat region around only MEKERK = ON.

When applied to a regulatory circuitry controlling self-renewal, CELLoGeNe uncovers the energy landscape containing attractors that correspond to experimentally observed stem cell states under various media. We managed to compress a multitude of experimental results into one framework, which provides further details like: (i) Assigned probabilities to possible stem cell states. (ii) Identified most likely destinations when perturbing a cell in a specific medium. (iii) Provided plausible explanations for leakage from pluripotency.

### Reprogramming MEF to iPSC

A natural application of CELLoGeNe is analysis of cell reprogramming systems. Here, we apply CELLoGeNe to a GRN governing the reprogramming of mouse embryonic fibroblasts (MEF) to induced pluripotent stem cells (iPSCs). Reprogramming experimental protocols have low conversion rates due to roadblocks preventing efficient reprogramming (***O’Malley et al., 2013; Chantzoura et al., 2015***). Using CELLoGeNe, we can analyse the full energy landscape and find unknown potential bottleneckss. We first applied CELLoGeNe to a GRN which has been updated from governing self-renewal and reprogramming in ***Dunn et al***. (***2019***). However, it was not possible to find any configuration of logical operators that fitted our data. We used experimental data from four stable states MEF, 2NG^−^, 3NG^−^, and iPSC, where cells in the 2NG^−^ and 3NG^−^ states tend stay in the same states without progressing towards iPSCs (***O’Malley et al., 2013***). After binarising the data we found that 2NG^−^ and 3NG^−^ correspond to the same cell state (***Figure 4C***). Since CELLoGene applied to the GRN in (***Dunn et al., 2019***) could not find the stable states from our data, we augmented the GRN with interactions found in literature concerning gene expression and perturbation experiments of reprogramming systems, see ***Figure 4A*** and ***Table 1***. Armed with our resulting network, we succeeded in finding configurations yielding valid energy landscapes.

**Table 1.** Components and sources and sources for the gene regulatory network governing reprogramming from MEF to iPSC.

| From | To | Effect | Reference(s) |
| --- | --- | --- | --- |
| LIF | Stat3 | activate | *Dunn et al. (2019); Niwa et al. (2009)* |
| Klf4 | Klf2 | activate | *Dunn et al. (2014, 2019)* |
| Esrrb | Klf2 | activate | *Xu et al. (2013, 2014); Yeo et al. (2014); Chen et al. (2008); Dunn et al. (2019)* |
| Tcf3 | Klf2 | repress | *Qiu et al. (2015)* |
| LIF | MEKERK | activate | *Graf et al. (2011)* |
| MEKERK | Tcf3 | repress | *Dunn et al. (2019)* |
| Klf4 | Tcf3 | activate | *Zhang et al. (2013); O'Malley et al. (2013)* |
| Stat3 | Gbx2 | activate | *Tai and Ying (2013); Dunn et al. (2019)* |
| Sox2 | Gbx2 | activate | *Xu et al. (2013, 2014)* |
| Oct4 | Gbx2 | repress | *Xu et al. (2013, 2014); Dunn et al. (2019)* |
| Stat3 | Klf4 | activate | *Dahéron et al. (2004); Bourillot and Savatier (2010); Dunn et al. (2019)* |
| Klf2 | Klf4 | activate | *Dunn et al. (2019)* |
| Gbx2 | Klf4 | activate | *Wang et al. (2017); Xu et al. (2013, 2014)* |
| Klf4 | Tbx3 | activate | *Dunn et al. (2019)* |
| Tcf3 | Tbx3 | repress | *Tam et al. (2008); Dunn et al. (2019)* |
| Oct4 | Tbx3 | repress | *Tam et al. (2008); Xu et al. (2013, 2014)* |
| Esrrb | Tbx3 | activate | *Adachi et al. (2018)* |
| Tfcp2l1 | Sall4 | activate | *Dunn et al. (2014); Tanimura et al. (2013); Dunn et al. (2019)* |
| Sox2 | Sall4 | activate | *Dunn et al. (2019)* |
| Tfcp2l1 | Esrrb | activate | *Dunn et al. (2014); Wang et al. (2019); Dunn et al. (2019)* |
| Nanog | Esrrb | activate | *Festuccia et al. (2012); Heurtier et al. (2019); Dunn et al. (2019)* |
| Sall4 | Esrrb | activate | *Tatetsu et al. (2016); Dunn et al. (2019)* |
| Tcf3 | Esrrb | repress | *Martello et al. (2012); Dunn et al. (2019)* |
| Sall4 | Oct4 | activate | *Tanimura et al. (2013); Dunn et al. (2019)* |
| Tcf3 | Oct4 | activate | *Ho et al. (2013); Dunn et al. (2019)* |
| Stat3 | Tfcp2l1 | activate | *Martello et al. (2013); Dunn et al. (2019)* |
| Esrrb | Tfcp2l1 | activate | *Wang et al. (2019); Dunn et al. (2019)* |
| Tcf3 | Tfcp2l1 | repress | *Qiu et al. (2015); Martello et al. (2012); Ye et al. (2017); Dunn et al. (2019)* |
| Oct4 | Tfcp2l1 | activate | *Martello et al. (2013); Stirparo et al. (2021)* |
| Oct4 | Nanog | activate | *Rodda et al. (2005); Boyer et al. (2005); Loh et al. (2006); Olariu et al. (2016); Dunn et al. (2019)* |
| Sall4 | Nanog | activate | *Zhang et al. (2006); Yang et al. (2008); Lim et al. (2008); Tatetsu et al. (2016); Dunn et al. (2019)* |
| Sox2 | Nanog | activate | *Rodda et al. (2005); Masui et al. (2007); Chen et al. (2008); Dunn et al. (2019)* |
| MEKERK | Nanog | repress | *Ying et al. (2008); Martello et al. (2013); Hamilton and Brickman (2014); Dunn et al. (2019)* |
| Tbx3 | Sox2 | activate | *Ivanova et al. (2006); Esmailpour and Huang (2012); Dunn et al. (2019)* |
| Nanog | Sox2 | activate | *Rodda et al. (2005); Boyer et al. (2005); Chickarmane et al. (2006, 2012); Dunn et al. (2019)* |
| Sall4 | Sox2 | activate | *Tanimura et al. (2013); Tatetsu et al. (2016); Dunn et al. (2019)* |
| Tcf3 | Sox2 | activate | *Yi et al. (2008); Dunn et al. (2019)* |
| MEKERK | Sox2 | repress | *Hamilton and Brickman (2014); Dunn et al. (2019)* |

**Figure 4.**
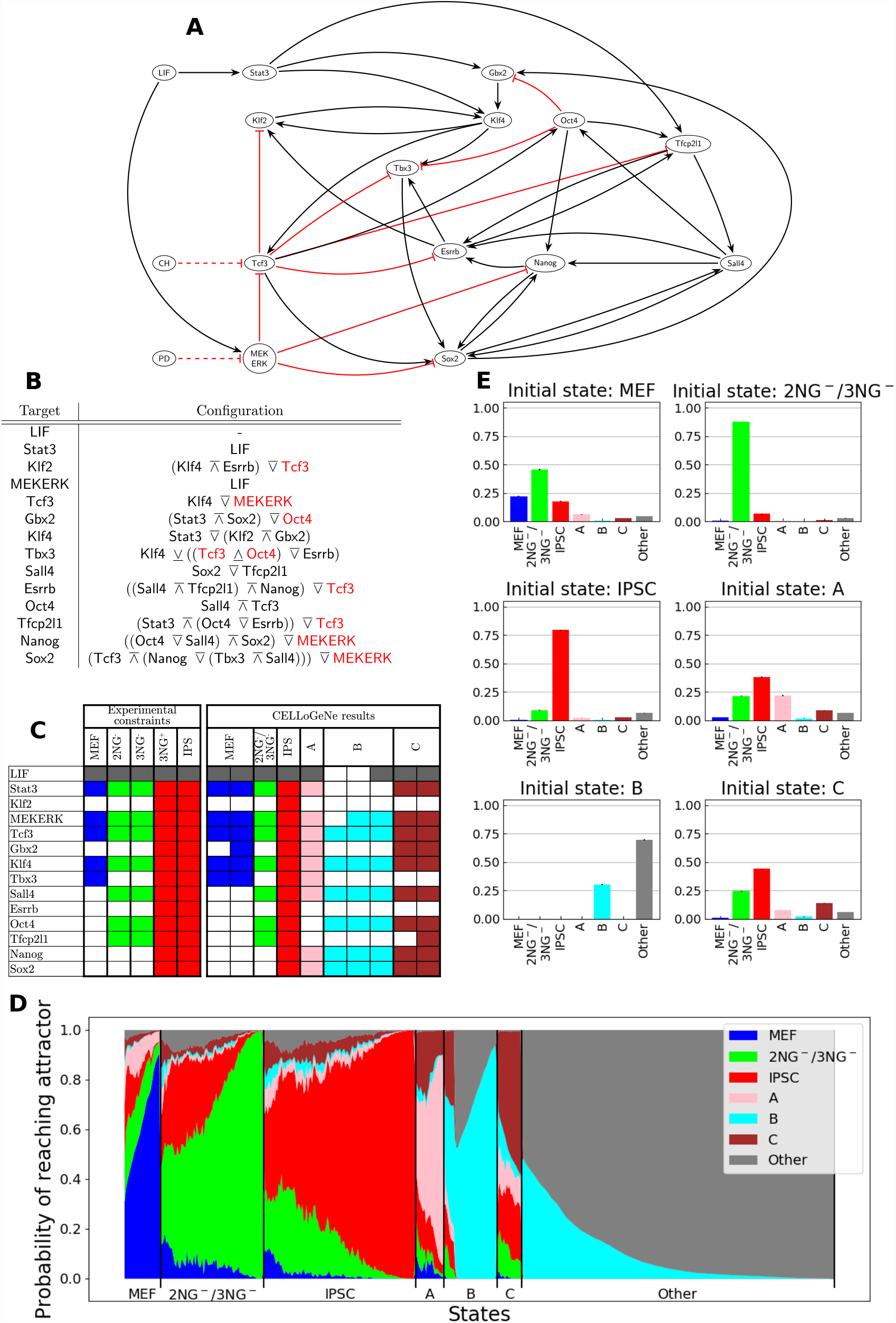
(**A**) GRN describing reprogramming of fibroblasts to iPSC. Black arrows represent activation, while blunted red arrows represent repression. **B**) Chosen configuration of logical operators. **C**) Experimental constraints (O’Malley et al, 2013) and attractors of the energy landscape. **D**) Overview of the relative basin sizes and strengths given from stochastic marble simulations. 1000 marbles per initial state were simulated with noise-level *β* = 1.5. **E**) Simulated reprogramming with each of the found attractors as initial states. The height of the bars represent the probability of ending up in an attractor. The marble simulations were run with *β* = 0.7 and 10 000 marbles per initial state. **Figure 4–Figure supplement 1**. The 2 valid configurations found when trying 10^6^ configurations. **Figure 4–Figure supplement 2**. Number of times the 10 most common attractors and the attractors in ***Figure 4C*** occur in the 10^6^ calculated energy landscapes. **Figure 4–Figure supplement 3**. Simulated reprogramming with different values on *β*.

Our GRN (***Figure 4A*** and ***Table 1***) consists of 14 nodes, 13 TFs and 1 medium component, LIF, yielding 2^14^ = 16 384 possible states. Since only LIF was used in our experiments (***O’Malley et al., 2013***), CH and PD inputs were not considered. Due to many input signals for each gene, it becomes computationally unfeasible to exhaustively test all configurations because of the prohibitively large number^5^ of possible configurations when using all 6 operators. Hence, we explored the configuration space stochastically, testing 10^6^ configurations, where a total of 2 valid configurations were found (***Figure 3–Figure Supplement 1***).

We found that the iPSC attractor is one of the 10 most prevalent minima in the 10^6^ calculated energy landscapes (***Figure 3–Figure Supplement 2***), occurring in 7.1 % of the landscapes^6^. Thus, the GRN is robust in its ability to reprogram to iPSCs with respect to different configurations of operators. The MEF and 2NG^−^/3NG^−^ attractors, on the other hand, only occur in 0.2 % and 0.7 % of the landscapes, respectively. Thus, the combined probability of randomly finding an energy landscape containing all three attractors is approximately one per million. This agrees with finding two valid configurations in the stochastic search. It is not surprising that MEF has a low probability of occurring since our GRN governs transition from the MEF state. However, the CELLoGeNe energy landscape needs to exhibit an attractor corresponding to MEF cells as this it the starting cell state in our experiments. This leads to important constrains on resulting operator configurations. We used a similar strategy as in the previous result section, constructing a manual configuration which we deemed to be biologically plausible (***Figure 4B***). This configuration yielded the required attractors, where MEF is a two-state degenerate attractor while 2NG^−^/3NG^−^ and iPS both are single state attractors (***Figure 4C***). Also three additional attractors were found, which we refer to as A, B and C (***Figure 4C***).

We performed marble simulations from each possible state without changing LIF from its initial value in order to probe the relative strengths and sizes of the basins of attraction (***Figure 4D***). We observed dominating attractors corresponding to MEF, 2NG^−^/3NG^−^ and iPSC states, with iPSC attractor being largest while MEF having smallest dominating region. We also observed smaller regions where attractors A, B and C are the dominating ones. This reveals that all the attractors’ basins have a region where they are the strongest, as opposed to the landscape for maintenance of pluripotency (***Figure 3E***). The fact that all of the three additional attractors A, B and C have regions where they dominate indicates that they all can act as bottlenecks. ***Figure 4D*** also shows that from around 40 % of the energy landscape states, i.e. the section furthest to the right, it is close to impossible to reach any of the experimentally known attractors. Attractor B and “Other” states are the only reachable attractors from this region. This region corresponds to those states where LIF is not present, showing that it is not possible to reach and maintain pluripotency without the adequate medium.

We conducted cell reprogramming simulations starting from the landscape’s six attractors (***Figure 4E***). When the cells were initialised in MEF, approximately a quarter of the cells stayed in MEF, barely half transitioned to 2NG^−^/3NG^−^ and a fifth transitioned directly to iPSC. The remaining cells ended up in A, C or other. Most of the cells initialised in 2NG^−^/3NG^−^ stayed there while the fraction of cells transitioning to iPSC was increased with higher noise level (***Figure 3–Figure Supplement 3***). When the cells were initialised in iPSC, most of the cells stayed in iPSC and a small fraction transitioned to 2NG^−^/3NG^−^. These three simulation results recapitulate experimental results observed in (***O’Malley et al., 2013***). The relative height of each bar varies with noise level (***Figure 3–Figure Supplement 3***), however, the general behaviour remains the same.

We performed a deeper analysis of the potential bottlenecks, by comparing the gene expressions for attractor states. Attractor A has the same genes ON as MEF, except three additional expressed genes: Sall4, Nanog and Sox2 (***Figure 4C***). Therefore, attractor A seems to be a roadblock when transitioning from MEF towards iPSC as more genes are turned ON. If the genes are turned ON in the wrong order, the cell risks to get stuck in attractor A. This could potentially happen if Sall4 is activated which could turn ON Sox2 and in turn activate Nanog, by following the logic of the operator configuration of the GRN (***Figure 4B*** and ***Figure 4A***). If, instead, the positive feedback loop with Oct4, Tfcp2𝗅1 and Sall4 is activated, then Oct4 could turn OFF Tbx3 and the cells get stuck in 2NG^−^/3NG^−^. If a few additional genes are also activated (Gbx2, Nanog and Sox2), then the cells get stuck in attractor C. Hence, it seems like attractor A mainly is a roadblock between MEF and iPSC, just like 2NG^−^/3NG^−^, while attractor C seems to be a roadblock between 2NG^−^/3NG^−^ and iPSC. Attractor B is not often reached from MEF or any of the other attractors, since it requires to turn OFF Stat3, which is directly activated by LIF. However, if attractor B is reached, it seems to be impossible for the cells to come back to the iPSC path.

When applied to a GRN governing reprogramming from MEF to iPSC, CELLoGeNe provides an overview of the energy landscape controlling cell fate decisions. With this broad overview, we identified known cell states as well as potential cell reprogramming bottlenecks. Marble simulations where we consider all possible states to be initial states gave a detailed overview of the possible roadblocks for reprogramming to iPS cells. This *in silico* analysis represents a powerful tool as huge number reprogramming starting points can be considered, which is virtually impossible experimentally. When we conducted simulations corresponding to performed reprogramming experiments, we obtained results which recapitulated experimental observations. Moreover, deeper analysis shed light on mechanisms that could lead to cells being trapped at the attractors corresponding to the observed and newly predicted reprogramming roadblocks.

## Discussion

In this study, we developed a novel framework, CELLoGeNe, which calculates energy landscapes for gene regulatory networks governing important processes like cell commitment, pluripotency maintenance and reprogramming. CELLoGeNe contains tools to analyse the resulting energy landscapes revealing cell states and reprogramming roadblocks through attractor identification and basins analysis. Getting access to the full energy landscape offers an overview of all the stable states of the biological system considered. This is not achievable with the standard methods of solving rate equations or updating a Boolean network. Considering a three-state logic, undesired symmetries are avoided compared to standard Boolean logic. CELLoGeNe is equipped with a tool to visualise multidimensional energy landscapes, giving direct insight about how the energy changes with gene expression variation. Cell lineage commitment and reprogramming to pluripotency along with its maintenance were investigated through the analysis of the basins of attraction surrounding stable cell states. We used a stochastic tool which provided a measure of relative strengths of the attractors and their basin’s extent, which was linked to the impact of reprogramming bottlenecks.

We applied CELLoGeNe to two systems describing aspects of maintaining and acquiring cell pluripotency. We analysed a GRN controlling self-renewal of pluripotency and found attractors in the energy landscape corresponding to experimentally observed stem cell states under various media. CELLoGeNe enabled us to assigned probabilities of finding the cells in a certain state. These probabilities were calculated by performing perturbation simulations of stable cell states in a specific medium and consequently identifying the most likely final cell state. When we applied CELLoGeNe to the core network in ***Dunn et al***. (***2014***) we identified more attractors than the ones experimentally and computationally presented in ***Dunn et al***. (***2014***). The discovery of these attractors offers a plausible explanation of the observed leakage from pluripotency (***Chambers et al., 2007; Chickarmane et al., 2012; Marucci, 2017***). CELLoGeNe was applied to a circuitry governing reprogramming from MEF to iPSC (***Takahashi and Yamanaka, 2006; Takahashi et al., 2007***). Cell reprogramming has a low efficiency and several experimental groups tried to improve this using various strategies: i) identifying new reprogramming factors (***Buganim et al., 2012; Shu et al., 2013***), ii) fine-tuning the levels of over expressed TFs (***Papapetrou et al., 2009; Radzisheuskaya et al., 2013***), iii) varying the order of introducing the factors used in reprogramming protocols (***Ho et al., 2013; Olariu et al., 2017a***), iv) monitoring reprogramming progress in a step-wise manner identifying bottlenecks (***O’Malley et al., 2013; Chantzoura et al., 2015***). In this study, we used CELLoGeNe for identifying new reprogramming bottlenecks, which can further be combined with experiments leading to improving reprogramming efficiency. To this end, we constructed a GRN controlling cell reprogramming and uncovered a corresponding energy landscape containing the MEF- and iPSC-states as well as the experimentally identified 2NG^−^/3NG^−^ roadblocks (***O’Malley et al., 2013***). The landscape contains three extra attractors which could correspond to potential roadblocks. The possibility for the cells to get stuck in these newly uncovered states links to observed inefficient conversion between MEF and iPSC. Analysing the gene expression defining these extra attractors sheds light on possible mechanisms leading to conversion inefficiency.

CELLoGeNe can be fused with an experimental CRISPR/Cas9-mediated genome wide knockout screen in reprogramming. The experiments could predict genes that can act as barriers of cell reprogramming. CELLoGeNe can be applied to GRNs containing the core reprogramming circuit augmented with the newly predicted genes. This would reveal the mechanisms through which the experimentally identified barriers act on preventing a successful cell reprogramming. Possible outcomes of adding the barrier genes to the circuit consist of emergence of new attractors that can act as a reprogramming roadblock or change of sizes of basins of attraction of existing attractors corresponding to experimentally identified roadblocks.

CELLoGeNe contains a tool that is capable of constructing a continuous energy landscape by interpolation, a feature which have not been used in this study. With a continuous energy landscape, it would be possible to construct an algorithm to map out optimal reprogramming paths which minimises the risk and prevents getting stuck at roadblocks. Furthermore, CELLoGeNe is a general tool which can be used on any developmental biological system. A strong candidate for CELLoGeNe applications is the T-cell development, as it provides an excellent model system for studying lineage commitment from a multipotent progenitor in general, because this biological system has been deeper experimentally and computationally analysed. In fact, even though CELLoGeNe was specifically developed for analysis of cell development and reprogramming systems, it can be applied to any system governed by a regulatory interaction network with states that can be binarised. To successfully apply CELLoGeNe to a non-biologically system, its network must have binary nodes affecting each other’s state either positively, negatively or neutrally.

We have developed a powerful framework for computing energy landscapes for GRNs enabling us to overview all possible stable cell states, which is inaccessible with other computational means. We applied CELLoGeNe to two biological systems achieving a better understanding of maintenance of pluripotency and offering a plausible explanation for spontaneous exit from multipotent stem cell fates. We also confirmed observed roadblocks and identified new potential cell reprogramming barriers, moreover, revealing their action mechanisms.

## Methods

### Binary representation of genes

The expression level of a gene can be normalised and represented with a number in the continuous interval [0, 1]. As an approximation, the gene expression can also be binarised into either being expressed (ON) or not expressed (OFF), which we will call a *logical gene*. Logical genes, thus, exist in the discrete space 𝔹 = {0, 1}. The *N* genes present in a GRN, i.e. the genes of interest, constitute the logical cell and exist in the space 𝔹^*n*^ = {0, 1}^*n*^, which we call the *expression* space. Since every gene can be either ON or OFF, the logical cell has 2^*n*^ states. Each state of a logical cell can be described by a vector ***s*** = (*g*_0_, *g*_1_, …, *g*_*n*−1_) of length *N* where each component represent a gene’s expression *g*_*i*_ ∈ 𝔹. Another useful representation is to convert the vector ***s*** into a binary number *s*. We construct *s* by letting *g*_*i*_ represent the *i*^th^ bit of the binary number. Formally, this is equivalent to 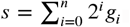. For instance, consider the state A=OFF, B=ON, C=ON, D=OFF and E=ON of the toy-network in ***Figure 2A***. The vector representation of this state is ***s*** = (0, 1, 1, 0, 1) which is equivalent to *s* = 10110_2_ = 22_10_ in base 2 and 10 respectively. Thus, each state ***s*** of the logical cell can be indexed uniquely by the integer *s* between 0 and 2^*n*^ − 1.

### Logical representation

With traditional Boolean logic, the activation A**→**B is equivalent to B = A, while the repression A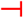B is equivalent to B = NOT A. However, as motivated in Methods-Three-state logic, this imposes an unwanted symmetry. Thus, we introduce a new three-state logic to describe network motifs.

The *expression* of the genes are kept in the 𝔹, however, we introduce a new space, the *effect* space 𝔼 = {−1, 0, 1}: negative, neutral and positive effect, where activation and repression are mapped differently. The effect an input gene’s expression has on a target gene is described with the mapping 𝔹 ↦ 𝔼 depending on the context: 0 always maps to 0 (0 ↦ 0), while 1 ↦ 1 when the input gene is an activator and 1 ↦ −1 when it is an inhibitor. The interpretations of the elements in the effect space are quite straightforward. If the effect is neutral, the expression of the target gene is not affected. If the effect is positive, the expression will become 1 if it was 0, or stay 1 if it was already ON. Similarly for negative effect, the expression will become 0. This can be be encapsulated with the forcing function

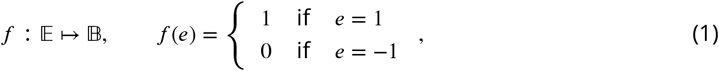

where *e* ∈ 𝔼 is the effect and *f* (0) is left undefined as 0 imposes no forcing.

### Combining input signals with operators

When multiple genes act as input for a target gene, their effect must be combined into a single input signal. In more mathematical terms, operators that map 𝔼 × 𝔼 ↦ 𝔼 are required.

A binary operator in ordinary Boolean logic has 2^2^ = 4 possible inputs, and is uniquely defined by the set of outputs it assigns to these; therefore, there are 2^4^ = 16 possible Boolean operators. Similarly, an operator in three-state logic has 3^2^ = 9 possible inputs, giving 3^9^ = 19683 possible operators. The vast majority of this prohibitively large number of operators are quite useless, so selection is needed.

We take *q*(*i, j*) to be the result when the operands are *i* and *j*, and define three properties that can be demanded of any reasonable operator:

- Idempotence — it is reasonable to assume that an interaction where all inputs are equal yields that same value as the output. Requires *q*(*i, i*) = *i*.
- Commutativity — there is no reason why the combination of genes should not be symmetric, and the lack of this property complicates the arithmetic. Requires *q*(*i, j*) = *q*(*j, i*).
- Associativity — also a basic arithmetic property. Requires *q*(*i, q*(*j, k*)) = *q* (*q*(*i, j*), *k*).

Placing these constraints on the standard Boolean operators leaves only AND and OR, which are usually the only ones that are included in Boolean models (along with unary NOT). On three-valued operators, the first two constraints leave only three free parametres of the original nine (for instance, *q*(0, −1), *q*(0, +1) and *q*(+1, −1)), on which the consequences of associativity can be worked out. For clarity, we can write *q* as a matrix **Q** such that *q*(*i* − 1, *j* − 1) = **Q**_*ij*_. With all three constraints, the result is a set of only nine operators, three of which are quite trivial:

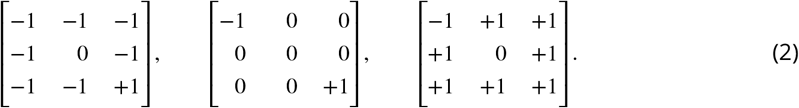

Except when required otherwise by idempotence, these just return a fixed value. The middle one is actually a rather straightforward generalisation of a two-valued Boolean operator: it acts as AND on {0, +1} and {0, −1}, and maps the additional input combination {±1, ∓1} to 0. The similarity to AND is shared with two more interesting operators:

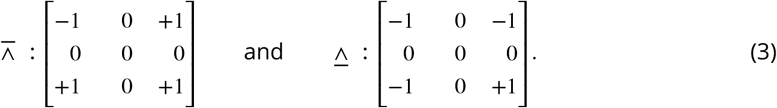

We have chosen symbols based on that for AND, ∧, with a bar above or below to symbolise mapping the different-sign inputs to +1 or −1, respectively. There exists a similar pair of operators for OR:

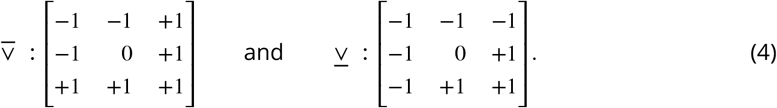

No associative analogue exists that maps different-sign inputs to 0. The two final operators are

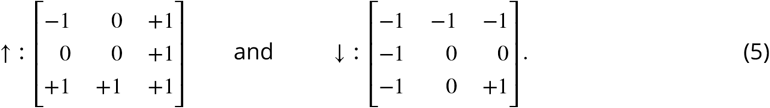

They return the maximum and minimum input value, respectively, and act as hybrids of AND and OR.

Either one of the pairs 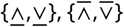 or {↓, ↑} provides an operator set that is fairly balanced with respect to the frequency of each output value. For greater flexibility, but also greater complexity, the full set of all six operators can be used.

By removing the requirement of idempotence, potentially useful operators such as xor can be added for Boolean logic. The corresponding modification gives 63 three-valued operators, none of which seems to be a good analogue.

### The discrete energy

Our goal is to build a stochastic model which shares the general behaviour of a deterministic model built on the same network. To do so in a straightforward way, we assign an energy *L* to each gene of a state whenever its expression status equals the result of its logical function, and an energy *H* whenever it does not. The energy contributions for each gene are added to form the total energy of a state. This is repeated for each state to form the energy of the entire network. If *L* < *H*, we thereby reward states that “obey the logic” with a favourable energy.

In more mathematical details, for each state ***s***, a gene g with expression value *a*_g_ ∈ 𝔹 is chosen. The other *n* − 1 genes serve as input to g and form the input space ***i***_g_ ∈ 𝔹^*n*−1^. To an input state ***i***_g_, there is a corresponding resulting effect *F* (***i***_g_) ∈ 𝔼 which is obtained by applying the combination operators, i.e. evaluating a logical function. The energy for state ***s***, *𝒯*(***s***), is calculated by adding the energy contributions from each gene *τ*(*a*_g_, ***i***_g_)

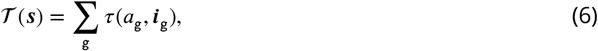

where

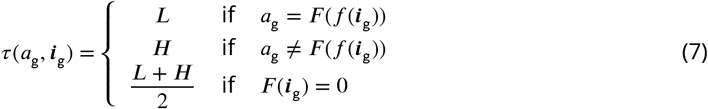

and *f* is defined in ***Equation 1***. Thus, the complete discrete energy landscape for the logical cell is represented by the vector **𝒯** where each component 𝒯_*s*_ is the energy 𝒯(***s***) of corresponding state ***s***. With *L* = −1, *H* = 1, a state with energy value 𝒯(***s***) = −*n* means that all gene expressions *a*_g_ agree with their corresponding forcing functions *F*(*f* (***i***_g_)) and vice versa for energy value 𝒯(***s***) = *n*.

### The continuous energy

The discrete energy is a function 𝒯 : {0, 1}^*n*^ → ℝ, which we can interpolate into a continuous energy *E* : [0, 1]^*n*^ → ℝ, which then allows each gene to have any degree of expression. Furthermore, there exists a useful form for *E*, in which it is first-order in all its variables, which makes the interpolation unique. Let ***s***_*i*_ be the expression of the *i*^th^ gene in the network, and ***s*** ∈ [0, 1]^*n*^ be the vector of all ***s***_*i*_. Then,

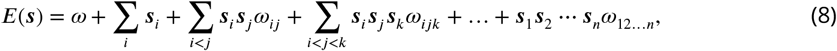

where all sums run between 1 and *N*. The *m*^th^ sum has 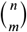 terms, so there are a total of

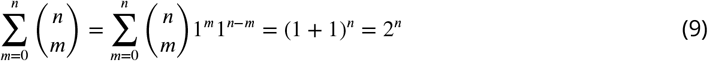

parameters *ω*. This is equal to the number of network states, so a bijection is guaranteed to exist between the discrete energy values and the parameters of the continuous energy.

This bijection has a simple explicit form. As mentioned above, each state is equivalent to a unique number in {0, 1, …, 2^*n*^ − 1}, so we use the vector 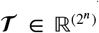 such that **𝒯**_*S*_ is the energy at the state ***s*** as described above. We then form the vector 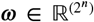 using the similar one-to-one correspondence ***ω***_*b*_ = *ω*_*ij*…_, where the binary representation of *b* has 1’s only at positions *i, j*, …. Using these vectors, we define the bijection as an invertible matrix **M** such that

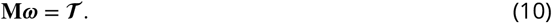

Assuming we have found such a matrix, the product

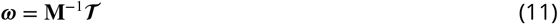

performs the transition from discrete to continuous energy. It can be proven that **M** and its inverse have the recursive Sierpiński triangle-like structure

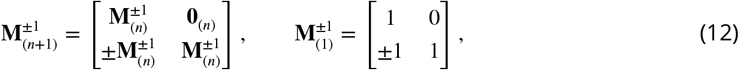

where **M**_(*n*)_ is the *N*-gene matrix of size 2^*N*^ × 2^*N*^, and **0**_(*N*)_ is the zero matrix of the same size.

There is a final benefit to this energy form. When it is paired with the entropy function

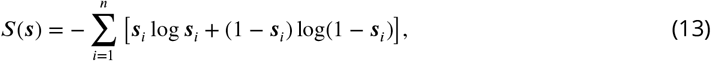

which contains no cross-terms or additional free parametres, we get a free energy function *F*(***s***) = *E*(***s***) − *T S*(***s***) (where *T* is the temperature) that generates sigmoid gain functions very closely approximating the Hill gain functions conventionally used in transcriptional dynamics (***Olariu et al., 2017a***). Thus, our model bridges the gap between stochastic and Boolean treatments.

### Three-state logic

In this section, we will give a motivation for the introduction of the three-state logic. We will start by considering the conventional Boolean approach and then extend it stepwise.

A basic Boolean model, which uses the operators AND, OR to build functions and NOT to represent repression, yields less than satisfactory energy functions when our method is applied. This can be demonstrated using the simple network B **→** A, which assigns A the function A = B. With *L* = −1, *H* = +1, and ***s*** = (A B), we get

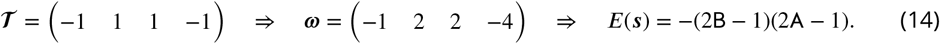

For B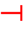A, which is represented by the function A = NOT B, we get the negative of this energy. The energy functions are entirely symmetric in B and A, even though the networks are directed.

The source of this unwanted symmetry is that B affects A equally much whether it is expressed or not. A perhaps more reasonable model is that a gene is only affected when its input is expressed; a gene with an inactive input is “free”, which we represent by giving it a third, “neutral” energy value between *L* and *H* (here, 0), regardless of the gene’s state. With these modifications, we get for B **→**A

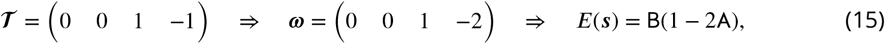

which properly reflects the directedness of the network. It also has an intuitive interpretation: the energy decreases when A is expressed, and the amount by which it does is proportional to B, i.e. how strongly A is activated. However, for B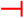A, we get

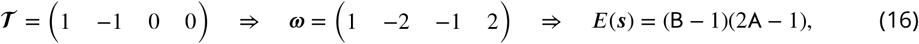

which is drastically different. It is directed, but goes against the above reasoning: B only affects A when it is not expressed.

To solve this, we stop using NOT altogether, and instead switch to the three-valued logic of true (1), false (0) and “negative true” (−1), the latter of which represents a repressing input, as described in Methods-Logical representation. A logical function whose result is −1 is obeyed if the targeted gene is not expressed, while 1 and 0 work as before. With the three-valued logic, B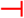A gives the same energy as ***Equation 15***, while B**→**A gives the negative of it, which is equally reasonable.

The three-valued logic results in presumably better energy functions, but it also incurs increased complexity, since AND and OR need to be replaced with the three-valued operators as described in Methods-Combining input signals with operators.

### Configurations of operators

When multiple genes act as input for a target gene, their effects need to be combined into a single input signal with an operator, as described in Methods-Combining input signals with operators. Which operator to use may be given from literature or experimental studies, but most often, it is not known which operator to use. Given that a real GRN most often has many nodes with several inputs, the network has many different configurations of operators where each configuration correspond to a potentially unique energy landscape. The total number of configurations of a network can be factorised into the product of the number of configurations there are for each target gene.

If there are *p* possible operators and a target gene has input from *k* genes, we denote the number of configurations for that gene *N*(*p, k*). For only one input gene, *N*(*p*, 1) = 1 trivially, since no inputs need to be combined. For only two input genes, *N*(*p*, 2) = *p*, since the only thing that can be changed is the operator that combines the two inputs. The rest is a matter of inductive reasoning, yielding the full recursive expression

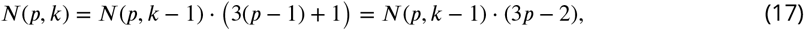

with the base cases given above. This has the closed form

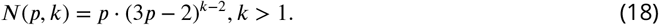

In ***Table 2*** some sample values for *N*(*p, k*) are presented, clearly demonstrating why we wish to limit the number of operators. Note that the values in the table are for one single target gene of the network only; to get the total number of the configurations of the network, the correct factors must be multiplied.

**Table 2.** Example values of *N*(*p, k*) for varying *k* (columns) and *p*. The case *k* = 2 case is left out, since *N*(*p*, 2) = *p*.

| $k$ | 3 | 4 | 5 | 6 | 10 | 15 | 20 |
| --- | --- | --- | --- | --- | --- | --- | --- |
| $p = 2$ | 8 | 32 | 128 | 512 | 131072 | 1.3e8 | 1.4e11 |
| $p = 3$ | 21 | 147 | 1029 | 7203 | 1.7e7 | 2.9e11 | 4.9e15 |
| $p = 4$ | 40 | 400 | 4000 | 40000 | 4.0e8 | 4.0e13 | 4.0e18 |
| $p = 6$ | 96 | 1536 | 24576 | 393216 | 2.6d10 | 2.7e16 | 2.8e22 |

### Testing configurations of operators

If more than one logical operator is used to combine gene input signals, there exist multiple configurations of operators. There are two main strategies on how to test different configurations: exhaustively test every combination, or randomly test a subset of combinations. For small GRNs or when few operators are used, an exhaustive search is suitable but, as understood from the previous section, sometimes the number of possible configurations is so large that a stochastic search must be used instead.

#### Exhaustive search

Given a set of genes and a set of operators, there is a large number of possible ways to combine these. All possible configurations are not distinct, thanks to commutativity and associativity. For instance, with {A, B, C} and {<u>∧</u>, <u>∨</u>}, there are eight configurations:

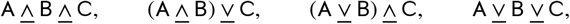

and all permutations of the genes in the middle two cases.

We now seek an optimal way to enumerate all non-redundant configurations, so that we can determine which one produces valid energy landscapes. Considering the example (A <u>∨</u> B) <u>∧</u> C, we note that if we change the first operator to <u>∧</u>, both (A <u>∧</u> B) and (…) <u>∧</u> C need to be recalculated. But if we change the second instead, we can reuse the result in parentheses. With this and other considerations, we can create specifications for an efficient method:

1. The outermost operators should be changed more often and operators inside parentheses should be changed only when all outer-level versions have been exhausted.
2. The same should apply to moving genes around within the parenthesis structure: more deeply nested genes should be accessed only after all permutations of outer genes have been used.
3. Changes should not be performed when commutativity and associativity make them superfluous. Such a method is devised below.

A convenient representation of configurations that allows for dealing with associativity and commutativity to avoid redundancy is binary trees: the leaves are genes, and the nodes are operators that combine their children. A *n*-gene compound can therefore be represented as a tree with *n* leaves, as illustrated in ***Figure 5A***. If each node “knows” its logical function, this representation enables reuse of functions in accordance with specifications 1 and 2. Each target gene in a GRN requires its own binary tree. For the trivial case where a gene only receives input signal from one other gene, the tree simply consists of one leaf. If a gene does not receive any input signal, e.g. a medium component such as LIF, its corresponding tree is empty.

**Figure 5.**
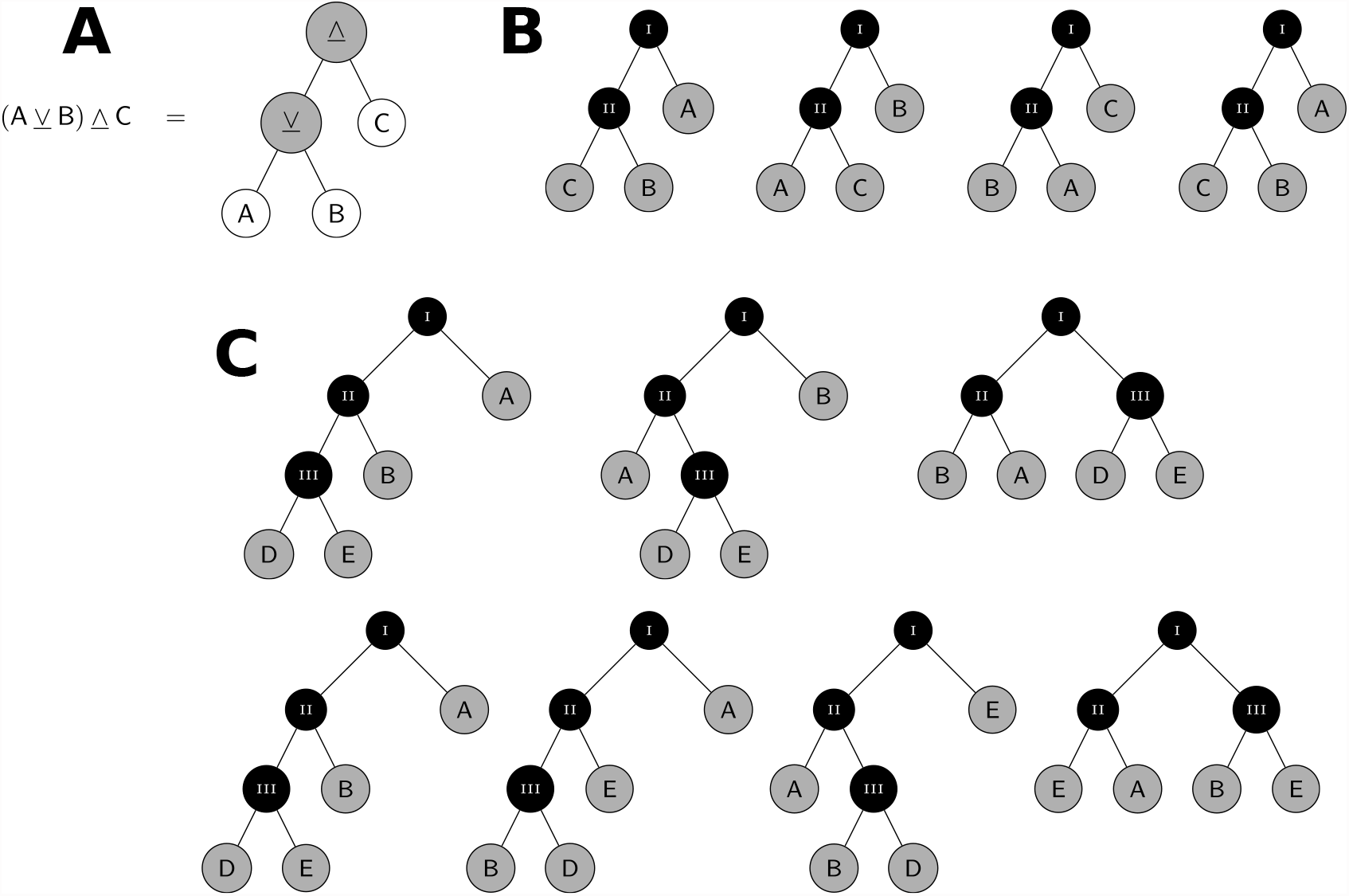
(**A**) A configuration of three genes represented as a binary tree. (**B**) Demonstration of the permutation of a tree. The black nodes are operator nodes, while the grey nodes can be either leaves or operator nodes with subtrees attached. (**C**) An extension of **B**, where node C is replaced with III and is revealed to have two children, D and E. In the top row, **B** is simply repeated. After that, a single permutation is performed around Node II, followed by the beginning of the next permutation cycle of node I, demonstrating how node E travels up the tree (node 4 will as well, later) and how the entire subtree with B and D is moved. **Figure 5–Figure supplement 1**. Flowchart describing the algorithm for trying all configurations of a gene. Applying it on the root once moves the tree to the next configuration. To save space, “op.” is used as an abbreviation of “operator”.

There is a recursive way to move around the parentheses in a tree-based manner that is well in line with the specifications. Since commutativity makes mirror images of a tree equivalent, we can without loss of generality only consider the cases where there are more leaves on the left side of the tree. The extreme case of this is when all operators are placed on the left edge of the tree, with only leaves on the right. This maximally left-heavy tree is the starting configuration of the method.

A demonstration of how it works is shown in ***Figure 5B*** on a minimal tree; after three permutations, it is back at the starting configuration. If node II has children of their own, the permutation procedure is recursively applied to it; this is shown in ***Figure 5C***. After a single such permutation, it is possible to make a fresh re-permutation around node I. When that is done, node II is allowed to take another step, and so on. At any level in the tree, a node only permits its child to permute one step when it has finished its own cycle of permutations.

This last rule, which we shall call the parent-first order, ensures two things. Firstly, as soon as the method reaches a node for which both children are leaves, it is guaranteed to be finished, since all levels above it have passed through all possible configurations. Secondly, as shown in ***Figure 5C***, a single permutation of the child replaces node B with a child of node C. It is then moved around by the parent, which puts it available for the parent’s parent (if present), etc. Applied inductively, this property guarantees that every leaf and every possible subtree moves around all possible positions in the tree, and thus the tree goes through all possible configurations regardless of its size.

The process of permuting a tree can be readily extended to include operator changes as well; the full procedure is given in ***Figure 5–Figure Supplement 1***. The algorithm becomes rather convoluted by the efforts to maximise function reuse and avoid redundant configurations—there are necessarily two associativity-safeguards labelled “same op. as left child?” in the figure—but stays implementation-friendly.

Operators are changed first because changing an operator is the cheapest possible change: it only requires the node’s own function to change (plus those of its parents), while a permutation also requires that of its left child to change. Also, a permutation changes which leaves are present in the child’s subtree, so the function needs to change its list of inputs in addition to its output; hence the “Update left child’s input” step.

Running this algorithm on the root of a tree until it hits FINISHED goes through all possible non-degenerate combinations of genes and operators. We note that larger trees are somewhat more costly to work with due to the larger number of deeply nested changes, so we should keep the trees sorted by ascending size when applying the algorithm.

Some further complications are straightforward to account for, should they be present for some reason:

- If the operators are non-commutative, insert “Update function with tree mirrored” after “Update function” in ***Figure 5–Figure Supplement 1***
- If the operators are non-associative, always answer “No” to “Same op. as left child?”

#### Stochastic search

Performing a stochastic search of the configuration space is substantially simpler than the exhaustive algorithm outlined above. For a given set of operators, a random binary tree representation (***Figure 5A***) is generated for each target gene in the GRN and the energy landscape is calculated. This is repeated until the desired number of configurations has been tested.

### Marble simulations

In order to measure the relative strengths of the attractors’ basins of attraction, we simulate cells performing weighted random walk in the energy landscape, analogous to letting marbles roll through the landscape until a minimum is reached. In each update, a cell can transition to a neighbouring state, i.e. changing one gene’s expression, or remain in the same state. The updates are repeated until a stopping criterion is met. By initialising a large instances of cells in every cell state and counting how many times the different attractors are reached as the final state, the probability of reaching each attractor from each cell state is obtained. By comparing the sizes of the regions of the landscape from which the attractors are reachable, and the probability of reaching each attractor from each cell state, the basins of attraction, and their relative strength, is mapped out.

The possible transitions for a cell state are weighted with the Boltzmann factor. A state *s* has energy *E*_*s*_ and the probability to transition to any of its *N* neighbours, or not make a transition, depends on the energy difference between the states (***Figure 6***). We define the transition probability from state *s* to *μ* as

**Figure 6.**
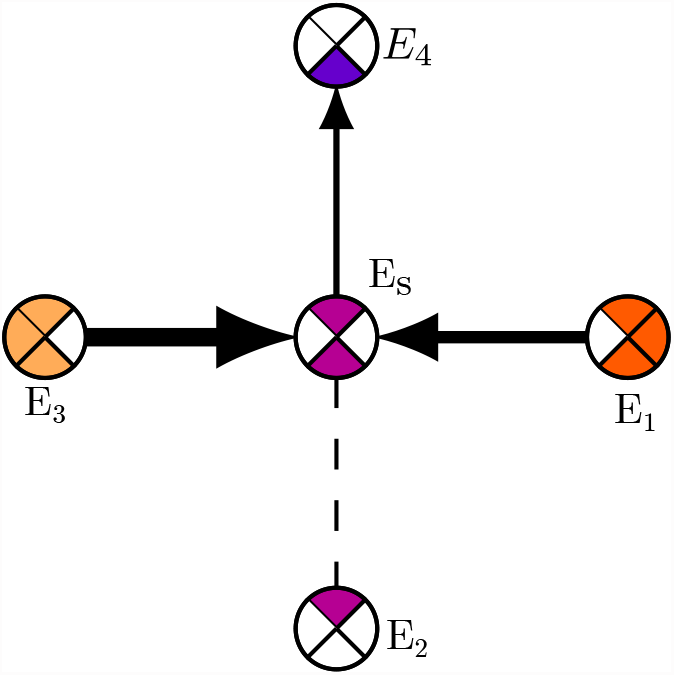
Example state *s* has energy *E*_*s*_ and its four neighbours have energies *E*_1…4_. Since state 4 is the only neighbour with energy *E*_4_ < *E*_*s*_ the transition *s* → 4 is the most probable, although all the other transitions are also possible due to the stochasticity. The probability is determined by the noise level *β* and energy differences between the states.

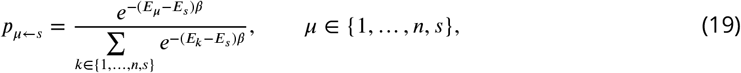

where *β* is the temperature parameter or noise level. By decreasing *β*, the noise level is increased and the probability to transition to a state with a higher energy increases. The transition to state *μ* is chosen as the smallest *μ* fulfilling

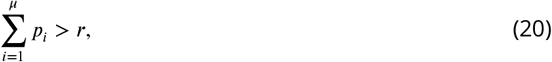

where *r* ∈ [0, 1) is a uniform random number. The cell state is updated until it has stayed in the same state during three consecutive updates, then it is considered to have reached a stable state.

### Visualisation of high-dimensional energy landscapes

The energy landscapes treated here are not particularly visualisation-friendly, since they exist in a space of high dimensionality. With traditional plotting methods, it is hard to depict of spaces in more than three dimensions. However, by utilising the the fact that an *N*-gene binary GRN has 2^*N*^ states, and each state has *N* neighbouring states^7^, we can construct somewhat reasonable representations of energy landscapes with up to seven dimensions. A known object which has 2^*N*^ states and *N* connections in *N* dimensions is a *N*-dimensional hypercube. By mapping the the corners of the energy function **𝒯** ∈ 𝔹^*N*^ to the corners of a hypercube, it just becomes a matter of plotting hypercubes onto two dimensions.

A 2-dimensional hypercube is simply a square. Thus, we let the four corners of a square represent the four states of a two-gene GRN (***Figure 7A***). We plot the square with rounded edges for future convenience. A three-dimensional hypercupe is a regular cube. In essence, a cube is just two squares vertically stacked with edges connecting corresponding corners. Hence, we can plot this in a flat version by arranging one square outside of another and connecting corresponding corners (***Figure 7A***). By the same principle, to represent a four-gene GRN, we can create a four-dimensional flat hypercube by connecting two flattened cubes (***Figure 7A***). By continuing this arrangement in a clever way, we can create flat hypercubes up to seven dimensions before they start becoming incomprehensibly cluttered (see ***Figure 3D*** for an exemplary energy landscape in seven dimensions). In principle, this plotting technique could, of course, be extended to arbitrarily many dimensions.

**Figure 7.**
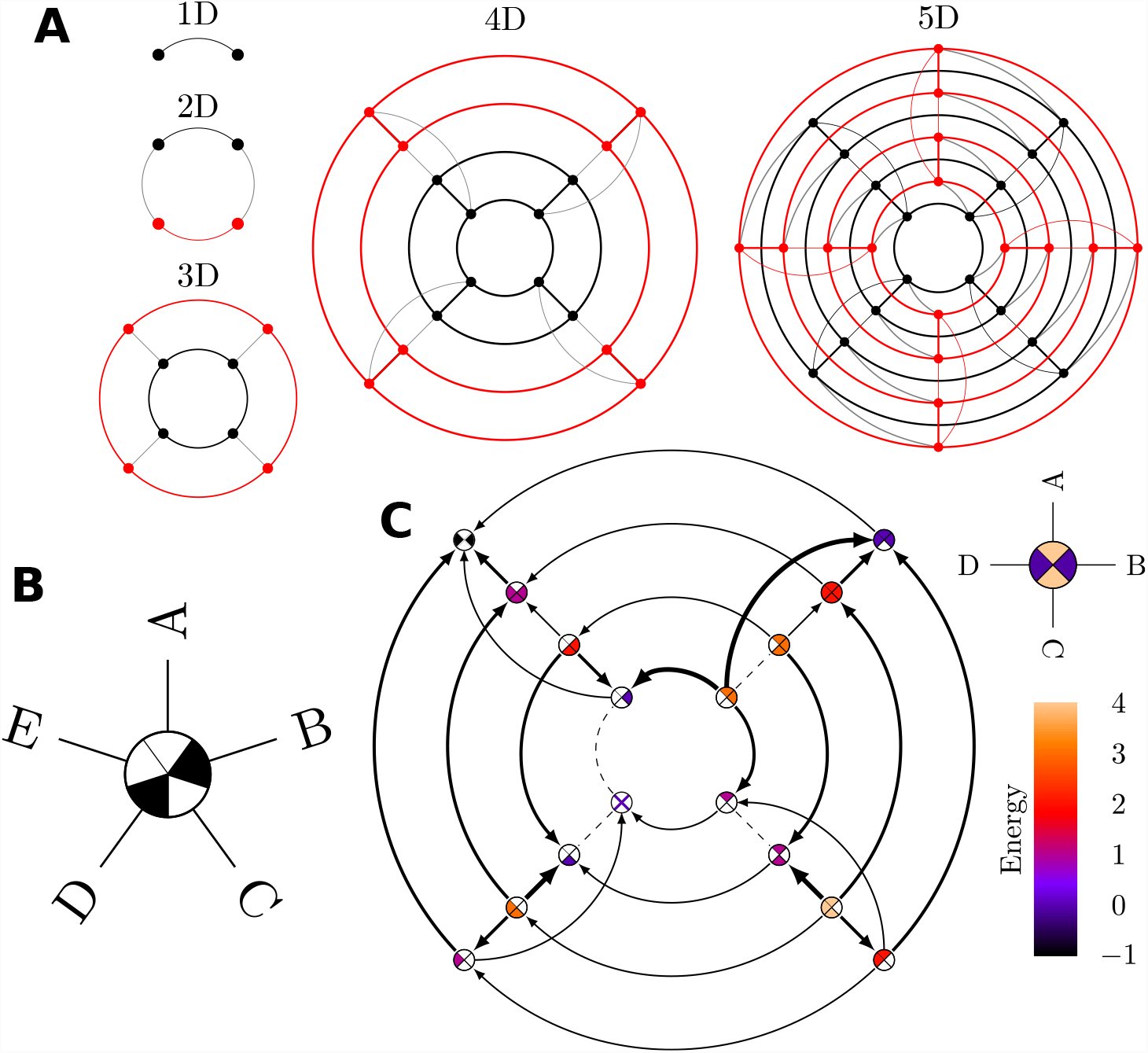
(**A**) Illustration of step-wise construction of an energy landscape in 1 to 5 dimensions. A landscape of dimension *N* consists of two instances of the landscape of dimension *N* − 1, one black and one red, which are connected with the grey lines. (**B**) Landscape node for a 5-dimensional energy landscape indicating the expression statuses of the 5 genes A to E, in this case A = OFF, B = ON, C = OFF, D = ON, E = OFF. (**C**) Example of a full energy landscape plot for a synthetic 4-dimensional landscape. Each landscape node represents a possible state of the logical cell where the colour of the node shows the energy value of that state, where the values are given by the colour bar to the right. Each node is divided into four sectors, where the sectors correspond to the genes as indicated in the legend above the colour bar. Filled sectors correspond to expressed genes while white sectors correspond to unexpressed genes. The arrows connect neighbouring states and points toward lower energies. The thickness of the arrows is a proportional to the magnitude of the energy gradient. Dashed lines illustrate flat segments in the landscape, i.e. two neighbouring states having the same energy value.

Now, we have a structure for the landscape plots in place, where each cell state maps to a corner on a hypercube. The next step is to add information. To clearly distinguish the states, we let each node in the graph be represented by a pie chart, where each piece of pie represents a gene. A gene is ON if its corresponding piece of pie is filled, and OFF if it is empty. For example, the state (A = OFF, B = ON, C = OFF, D = ON, E = OFF) is depicted as shown in ***Figure 7B***. The energy value of each state is represented by a colour which maps to a colour bar. The final property of the basic version of the landscape plots is that every edge describes the energy difference between the connected states. The edge is drawn as an arrow, pointing towards the state with lower energy, and its width is proportional to the magnitude of the energy difference. Neighbouring states with the same energy are connected with a dashed line. An example of a complete energy landscape is displayed in ***Figure 7C***.

With this plotting technique, it is, as stated above, possible to represent up to seven-gene networks before the plots get prohibitively large. However, there are ways to circumvent this obstacle. One possible trick is to only plot hyperplanes of interest. Then, one chooses up to seven genes of interest to be displayed, and the remaining genes are fixed. Another trick is to merge several genes that behave similarly in the system of interest into one single dimension. If this can be done with several groups of genes, this has the possibility to reduce the dimensionality substantially. The downside of this trick is that states that are not neighbours in the full energy landscape will appear as neighbours in the reduced landscape. However, this can still be useful in some circumstances.

More details can be added for further clarity. Attractor states and their edges can be colour coded. For full landscapes, the basins of attraction may be illustrated by a coloured ring around the landscape node. The ring is partitioned and colour coded according to which attractors’ basins the state is part of. This requires that the attractors are colour coded as well. These additional details are utilised in ***Figure 2D***.

## Acknowledgments

The authors would like to thank Carsten Peterson and Bo Söderberg for discussions at various stages of the project. VO gratefully acknowledges the support of the US National Institutes of Health (USPHS grant R01HL119102) and Crafoordska Stiftelsen.

## Author contribution

EA, MS and VO designed the study. EA and MS built the CELLoGeNe software platform. EA conducted the biological applications and computational analysis. KK provided, expertly curated and binarised the experimental data. KK explicated and conceptualised the use of experimental data. EA, MS and VO wrote the manuscript. All authors provided inputs and comments on the manuscript.

---

**Figure 3–Figure supplement 1.**
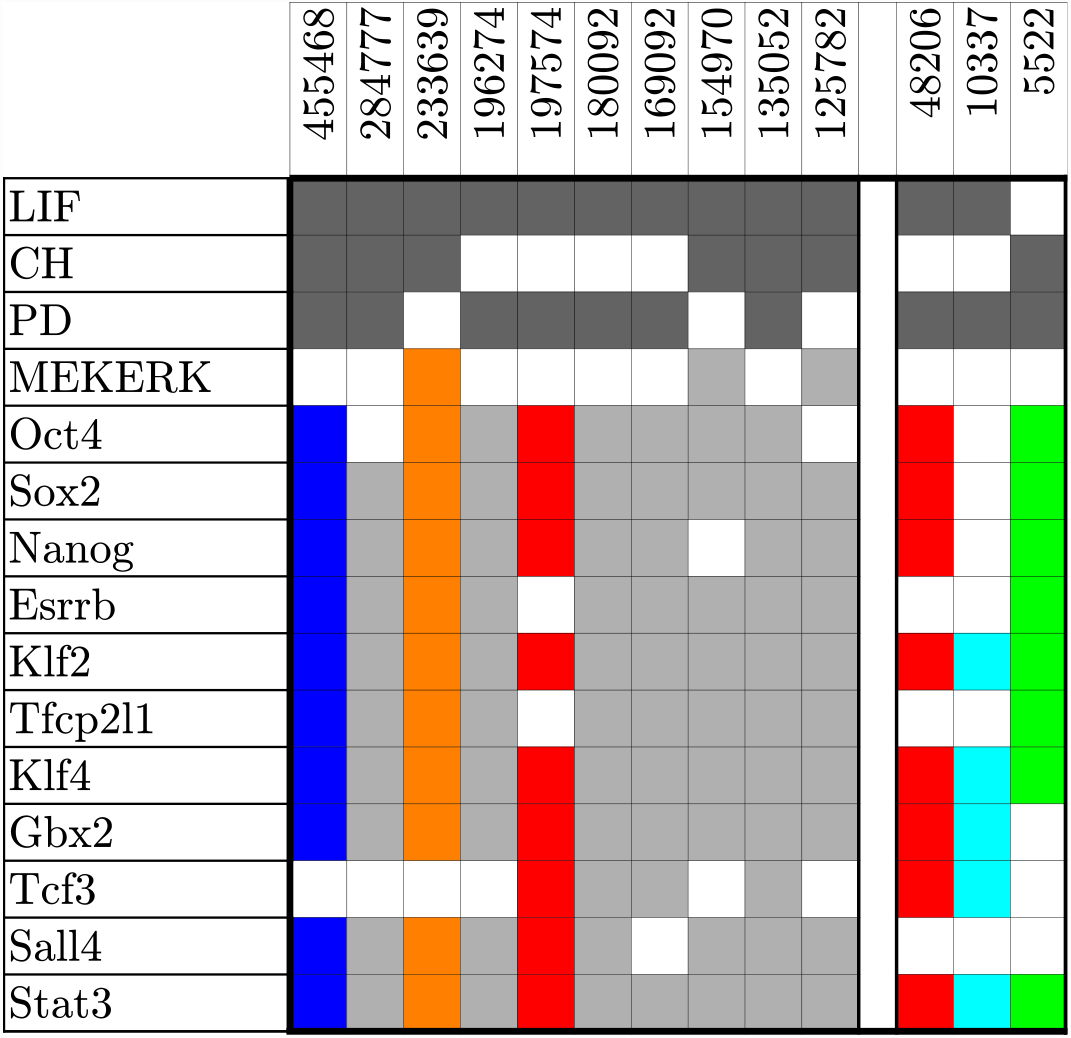
Number of times the 10 most common attractors and experimental constraints occur in the 10^6^ calculated energy landscapes.

**Figure 3–Figure supplement 2.**
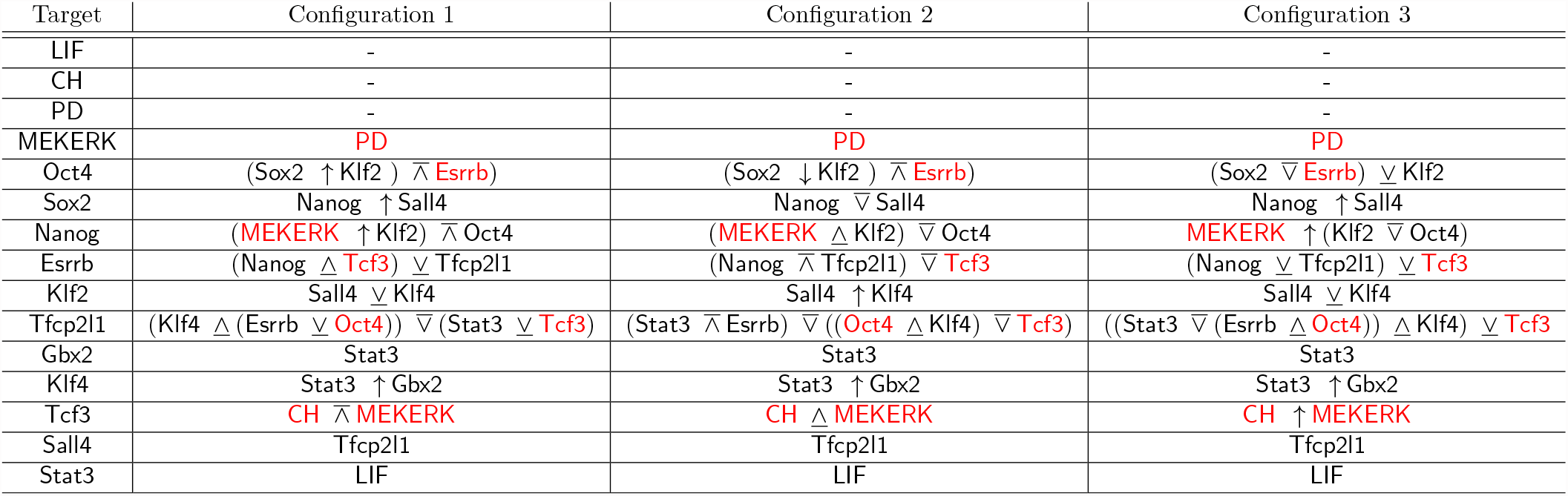
The 3 configurations with the lowest degeneracy out of the 163 valid configurations found when trying 10^6^ configurations.

**Figure 4–Figure supplement 1.**
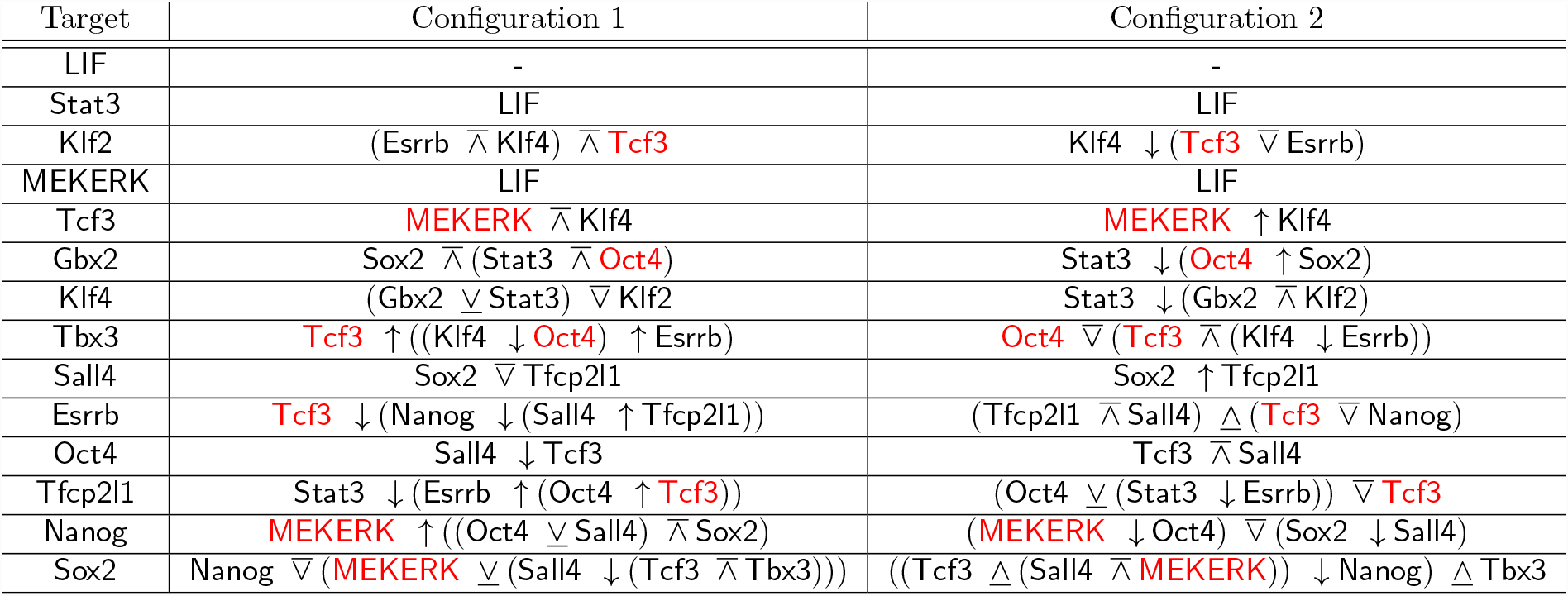
The 2 valid configurations found when trying 10^6^ configurations.

**Figure 4–Figure supplement 2.**
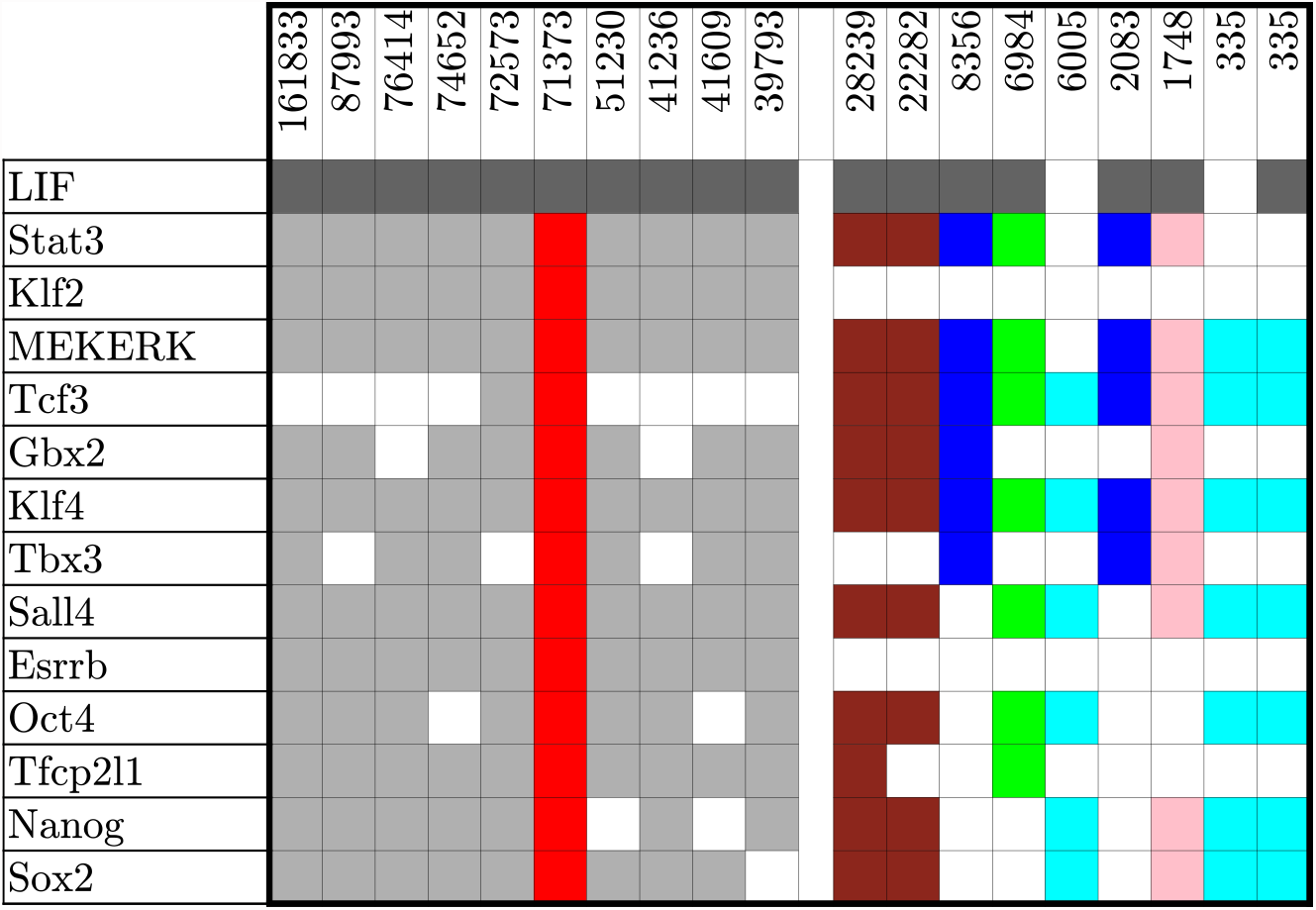
Number of times the 10 most common attractors and the attractors in **Figure 4C** occur in the 10^6^ calculated energy landscapes.

**Figure 4–Figure supplement 3.**
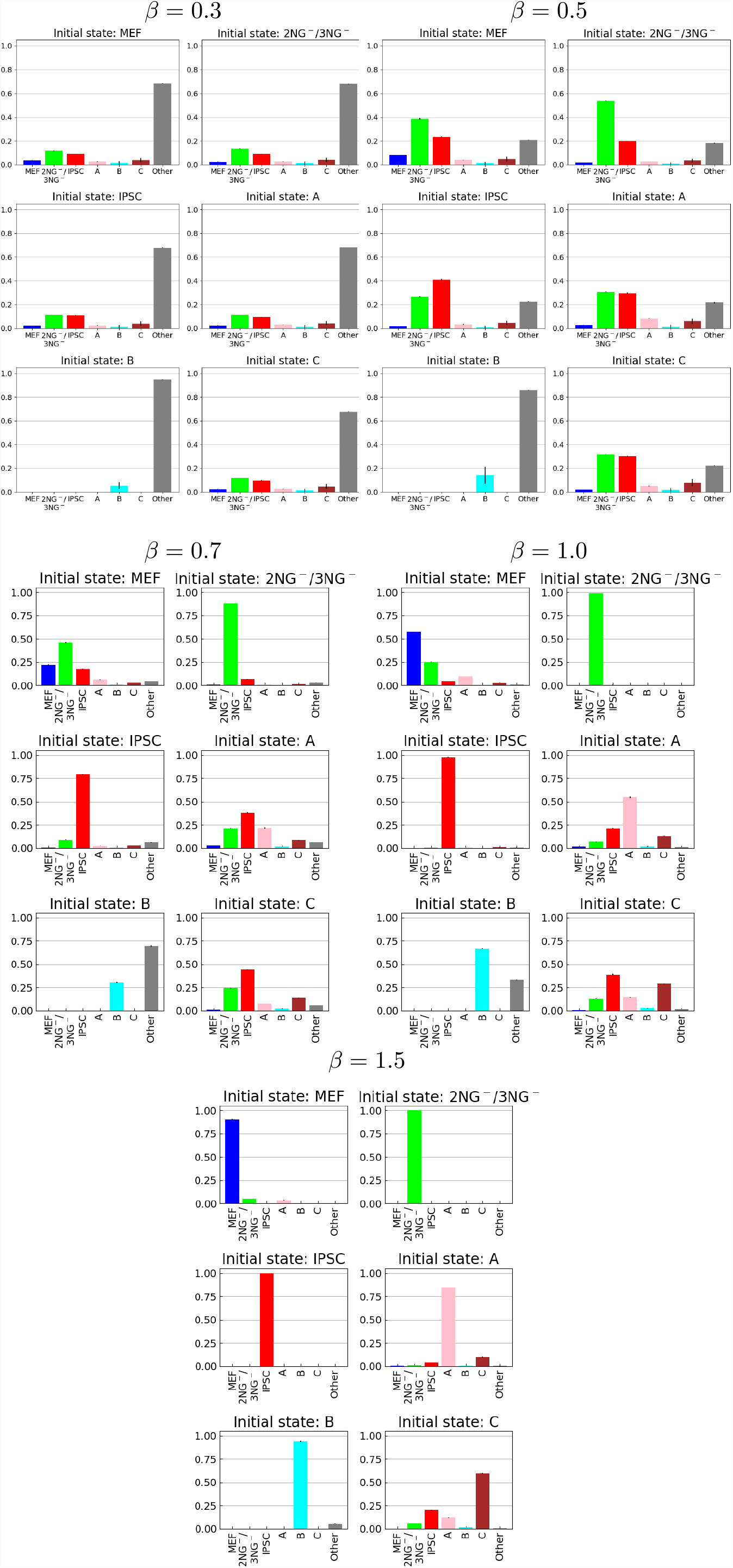
Simulated reprogramming with different values on *β*.

**Figure 5–Figure supplement 1.**
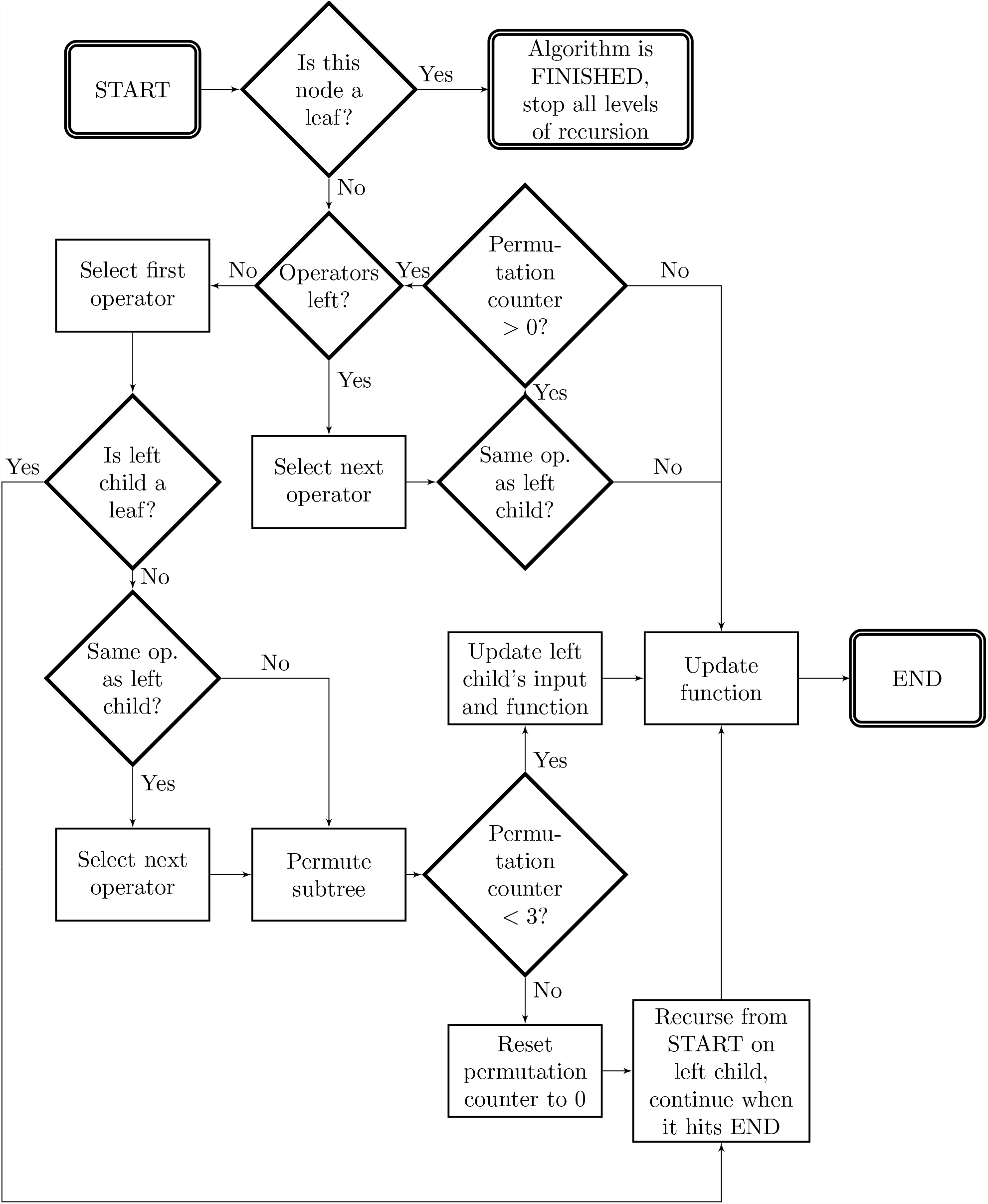
Flowchart describing the algorithm for trying all configurations of a gene. Applying it on the root once moves the tree to the next configuration. To save space, “op.” is used as an abbreviation of “operator”.

## Footnotes

1 The exact number of possible configurations is 28 179 280 429 056 ∼ 10^13^.

2 In order to not get an incomprehensibly large plot, we merged groups of genes into shared dimensions as denoted by the legend to the right in ***Figure 3D***.

3 The pluripotent genes are here considered to be Oct4, Sox2, Nanog, and Klf 2.

4 Neighbouring cell states have Hamming distance 1, e.g. the states 10010 and 10011 in a 5-gene system.

5 The exact number is 26 142 282 979 403 407 520 956 416 (∼ 10^25^).

6 As reference, the most prevalent minimum occurs in approximately 16 % of the energy landscapes.

7 With neighbouring state we mean states where only one bit changes, i.e. states with Hamming distance equal to 1.

